# Comprehensive Analysis of Two *Shank3* and the Cacna1c Mouse Models of Autism Spectrum Disorder

**DOI:** 10.1101/068866

**Authors:** Patricia Kabitzke, Daniela Brunner, Dansha He, Pamela A. Fazio, Kimberly Cox, Jane Sutphen, Lucinda Thiede, Emily Sabath, Taleen Hanania, Vadim Alexandrov, Randall Rasmusson, Will Spooren, Anirvan Ghosh, Pamela Feliciano, Barbara Biemans, Marta Benedetti, Alice Luo Clayton

**Affiliations:** PsychoGenics, Inc., Tarrytown, NY, USA; Department of Psychiatry, Columbia University, New York, NY, USA; Department of Physiology and Biophysics, SUNY Buffalo School of Medicine and Biomedical Sciences, Buffalo, NY, USA; Roche Pharma Research and Early Development, NORD, Roche Innovation Center, Basel, Switzerland; Simons Foundation Autism Research Initiative, New York, NY, USA

## Abstract

To expand, analyze and extend published behavioral phenotypes relevant to autism spectrum disorder (ASD), we present a study of three ASD genetic mouse models: Feng’s *Shank3^tm2Gfng^* model, hereafter *Shank3/F*, Jiang’s *Shank3^tm1Yhj^* model, hereafter *Shank3/J*, and the Cacna1c deletion model. The *Shank3/F* and *Shank3/J* models mimick gene mutations associated with Phelan-Mcdermid syndrome and the Cacna1c model recapitulates the deletion underlying Timothy syndrome. The current study utilizes both standard and novel, computer-vision based behavioral tests, the same methdology used in our previously published companion report on the *Cntnap2* null and 16p11.2 deletion models. Overall, some but not all behaviors replicated published findings. Those that replicated, such as social behavior and overgrooming in *Shank3* models, also tended to be milder than previous reports. The *Shank3/F* model, and to a much lesser extent, the *Shank3/J* and Cacna1c models, showed hypoactivity and a general anxiety-like behavior triggered by external stimuli which pervaded social interactions. We did not detect deficits in a cognitive procedural learning test nor did we observe perseverative behavior in these models. We did, however, find differences in exploratory patterns of Cacna1c mutant mice suggestive of a behavioral effect in a social setting. In addition, *Shank3/F* but not *Shank3/J* KO or Cacna1c HET showed differences in sensory-gating. Discrepancies in our current results from previous reports may be dependent on subtle differences in testing conditions, housing enrichment, or background strain. Both positive and negative results from this study will be useful in identifying the most robust and replicable behavioral signatures within and across mouse models of autism. Understanding these phenotypes may shed light of which features to study when screening compounds for potential therapeutic interventions.

## Introduction

Several mouse mutants have been created based on evidence derived from human genetic studies, implicating copy number and single nucleotide variants in autism spectrum disorder (ASD). These models offer an opportunity to advance drug development in ASD based on the human condition. Therefore, we embarked on a large effort to phenotype five ASD mouse models utilizing comparable assays to find those which may be robust endpoints amenable to future drug screening. We provide back-to-back characterization of five different models of ASD through our previous companion paper on the first two models, *Cntnap2* and 16p11 deletion, [1] and present here the results pertaining to the third, fourth, and fifth models, namely, two distinct *Shank3* knockout (KO) models and the Cacna1c heterozygous (HET) model. We chose these models because of their strong construct validity based on human genetic evidence and because of their widespread use in the scientific community due to their availability through The Jackson’s Laboratories as part of the Simons Foundation Autism Repository. In addition, considerable effort has been placed in the backcrossing of all models to the C57 background, whenever possible, to enable comparisons across models. We chose a broad battery of behavioral endpoints targeting both core and auxiliary domains of ASD. Although we readily acknowledge the potential caveats of mouse behavior in homology and translatability to humans with ASD, we believe that the analysis of complex function in mice provides valuable information regarding downstream effects of the chosen genetic manipulations. Thus, we focused our phenotyping efforts on 1) tests that had been used to characterize the models in the initial phenotyping publications, 2) tests that provide information on the models’ social, perseverative, cognitive, developmental and motor functions, and 3) unbiased, data-driven phenotyping tests to uncover unexpected phenotypes. We present, in these two papers, the complete set of results with the hope that researchers can use the data to further ASD discovery science and also for potential screening of therapeutics. The third and final paper in the series will present a replication of our main findings using separate cohorts of *Cntnap2* and *Shank3* KO models as well as bioinformatics analyses comparing all five models to each other.

The relevance of the *Shank3* gene for human disease became apparent once the 22q13.3 deletion, or Phelan-McDermid syndrome, which includes its coding sequence, was shown to be associated with a syndrome comprising hypotonia and developmental delay, intellectual disability in association with severe language delay and autistic-like behavior [2].

The postsynaptic density (PSD) is a synaptic area with a dense concentration of proteins involved in fast signal transmission. Proteins of the excitatory PSD, like Shank, have various functions, from signal transduction, structural regulation, to metabolism. Shank domains include an N-terminal ankyrin repeat, a SH3, and a PDZ domain in addition to a proline-rich sequence (with binding sites for Homer and cortactin) and a C-terminal SAM domain (important for synaptic localization), in keeping with their ability to form complexes with receptors, and with signaling and cytoskeletal proteins [3, 4]. The complexity of the Shank family of proteins is further increased by the existence of 3 genes and isoforms resulting from alternative splicing.

All SHANK family proteins (1, 2, and 3) have been associated with idiopathic ASD and linked to synaptic dysfunction [5]. *De novo* or inherited *Shank3* mutations associated with ASD affect its localization at the tip of the F-actin fibers, which disrupt spine formation and morphology [6].

Several deletions of the *Shank3* gene have been accomplished in mice. One of the models created referred to as *Shank3B* KO in Peça et al. [7] harbors a deletion of exons 13-16 of the PDZ domains (*Shank3^tm2Gfng^* or *Shank3/F* hereafter). Jiang and colleagues used gene-targeting techniques to delete exons 4–9 of the *Shank3* gene (*Shank3^tm1Yhj^* or *Shank3/F* hereafter) resulting in the absence of the *Shank3α* and *β* isoforms [8]. Timothy syndrome, caused by a deletion of Cacna1c, is associated with morphological characteristics such as webbed fingers and toes and facial dimorphism, and is associated with heart abnormalities and autism [9]. The mutations change the configuration of this important ion channel and result in increased calcium influx. Homologous mutations in mice proved lethal although viability was restored with the insertion of a neo cassette, and thus the line was termed TS2-neo in the original phenotypic report [10].

We present here an assessment of these two *Shank3* murine models of ASD and of the Cacna1c Timothy model using a comprehensive behavioral battery. This test battery was purposely designed to broadly examine multiple functional domains in an effort to identify robust, replicable behavioral phenotypes regardless of whether those behaviors are anthropomorphically considered autistic-like. As such, the battery included, but was not restricted to tests of the “autistic triad” (social impairment, communication deficits, repetitive behaviors), and also incorporated proprietary platforms to discover possible unexpected phenotypes to complement standard tests.

## Results

### SmartCube

As previously described in detail [1], SmartCube is a high-throughput automated behavioral platform that allows rapid, comprehensive phenotyping of mutant models and detection of behavior patterns influenced by drugs or any other manipulation. It employs computer vision and mechanical actuators to detect spontaneous behavior as well as evoked ones elicited by anxiogenic and startling stimuli. Behavioral readouts include locomotion, trajectory complexity, body posture and shape, simple behaviors and behavioral sequences [11-13]. Supervised machine learning algorithms comb the set of more than 1400 collected features (see S1 Methods). Data are plotted in the 2D feature space that best separates mutant from wild type groups, and the overlap between the two datasets is used as a discrimination index. To assess the robustness of the separation, the machine-learning algorithm is trained and tested many times with different data subsets using correct and randomized labels. The overlap between the resulting distributions of discrimination indexes is used to estimate the probability that the results could be due to chance.

The discrimination index estimates the differences between two groups when all features are considered as a whole. For *Shank3/F* WT and KO mice, this value reached 74% (Figs 1A and 1B). The probability that this discrimination value could be obtained by chance, in a similar dataset, is *p*< 0.02. Top features contributing to this difference were reduced speed, sniffing, rearing, locomotion and increased freezing, latency to approach a stimulus and grooming in the KO compared to WT mice (Figs 2; S1 Table). In contrast, the discrimination between the *Shank3/J* WT and KO mice reached 66%. The probability that this discrimination value could be obtained by chance, in a similar dataset, is *p*< 0.09 (Figs 1C and 1D). The top feature was increased freezing (Fig 2; S1 Table). For Cacna1c HET and WT mice, this value reached 88% (Figs 1E and 1F). The probability that this discrimination value could be obtained by chance, in a similar dataset, is *p*< 0.001. Top features, those behaviors that contribute the most to the separation between the two genotype groups, were fewer abrupt movements, locomotion bursts, and scanning and increased freezing in the HET compared to WT mice (Fig 2; S1 Table). To provide a comparison across models, we represent the same features for the three lines, even if differences were not found for all three models. It is important to note that the aggregate of many small subtle behavioral changes could contribute to an overall robust signature in SmartCube.

**Fig 1.**
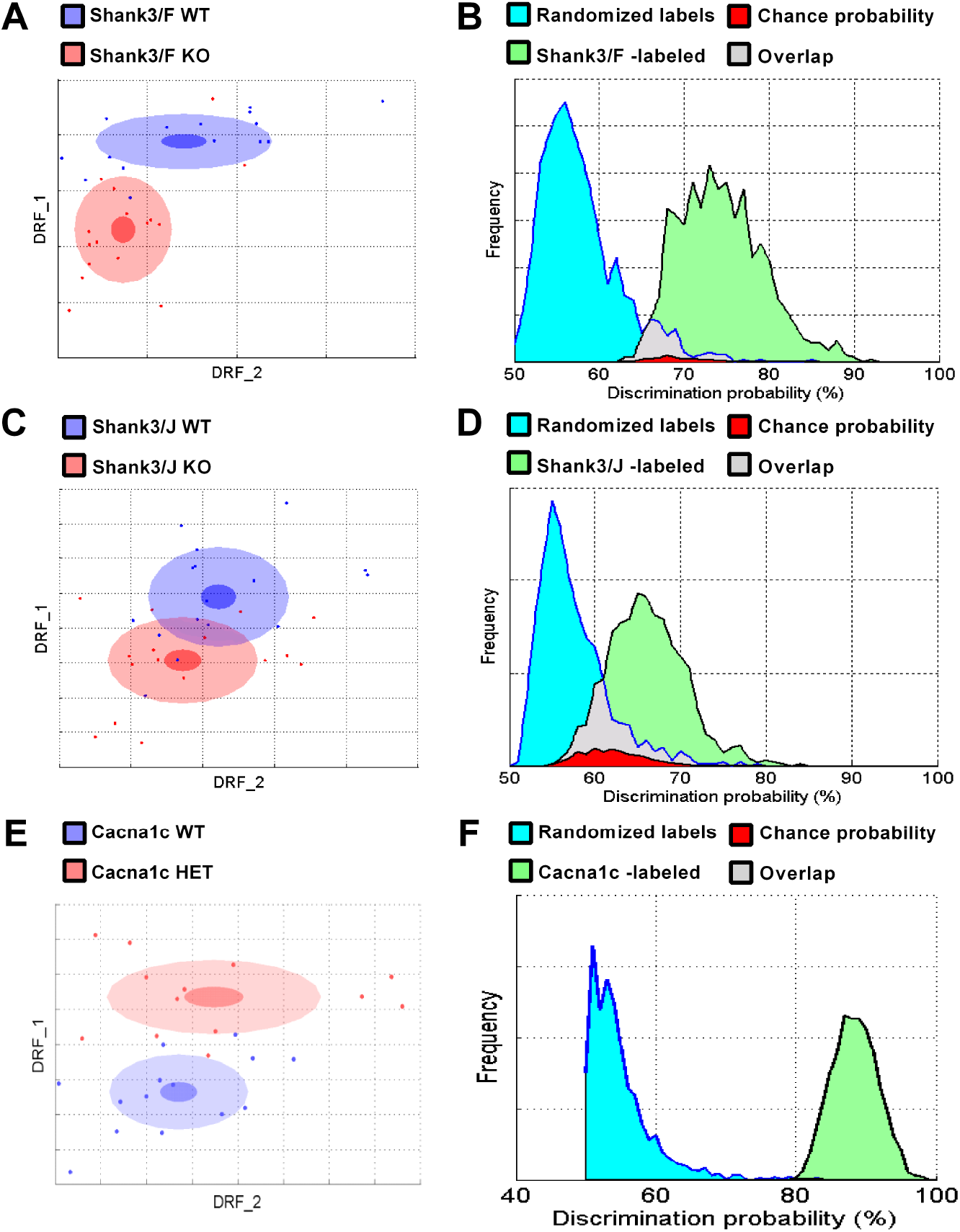
SmartCube found different degrees of separation between mutant Shank3/F, Shank3/J mice, and Cacna1c mice as compared to their corresponding WT control littermates. A & C: As described previously in detail [1], to build a 2D representation of the multidimensional space in which the two groups are best separated, we first find statistically independent combinations of the original features, pick the two new composite features that best discriminate between the two groups, and used them as x- and y-axes (drf 1 and 2; see S1 Methods). Each dot represents either a WT (blue) or a mutant (red) mouse. The center, small and large ellipses are the mean, standard error and standard deviation of the composite features for each group. The overlap between the groups is used to calculate the discrimination index, which measures how reliably a classifier can be trained to discriminate between the two groups (the more overlap, the worse the discrimination). B & D: To estimate how likely it is that such separation is simply due to chance, the obtained classifier is challenged many times with correctly labeled samples (green distribution) or with randomized labels (blue distribution). The overlap between these two distributions (in red) represents the probability of obtaining the observed discrimination by chance. A: The Shank3/F model separates well from the WT group. Note the spread and position variability of the individual mice. B: 74% discrimination between Shank3/F and WT mice could be found by chance with p< 0.02. C: The Shank3/J standard deviation ellipse overlaps considerably with the WT control ellipse. D: A 66% discrimination index for the Shank3/J versus WT comparison can be found by chance with p< 0.09. E: The Cacna1c model separates well from the WT group. Note the spread and position variability of the individual mice. B: 88% discrimination between Cacna1c and WT mice could be found by chance with p< 0.001. n=13-16 mice per genotype/line.

**Fig 2.**
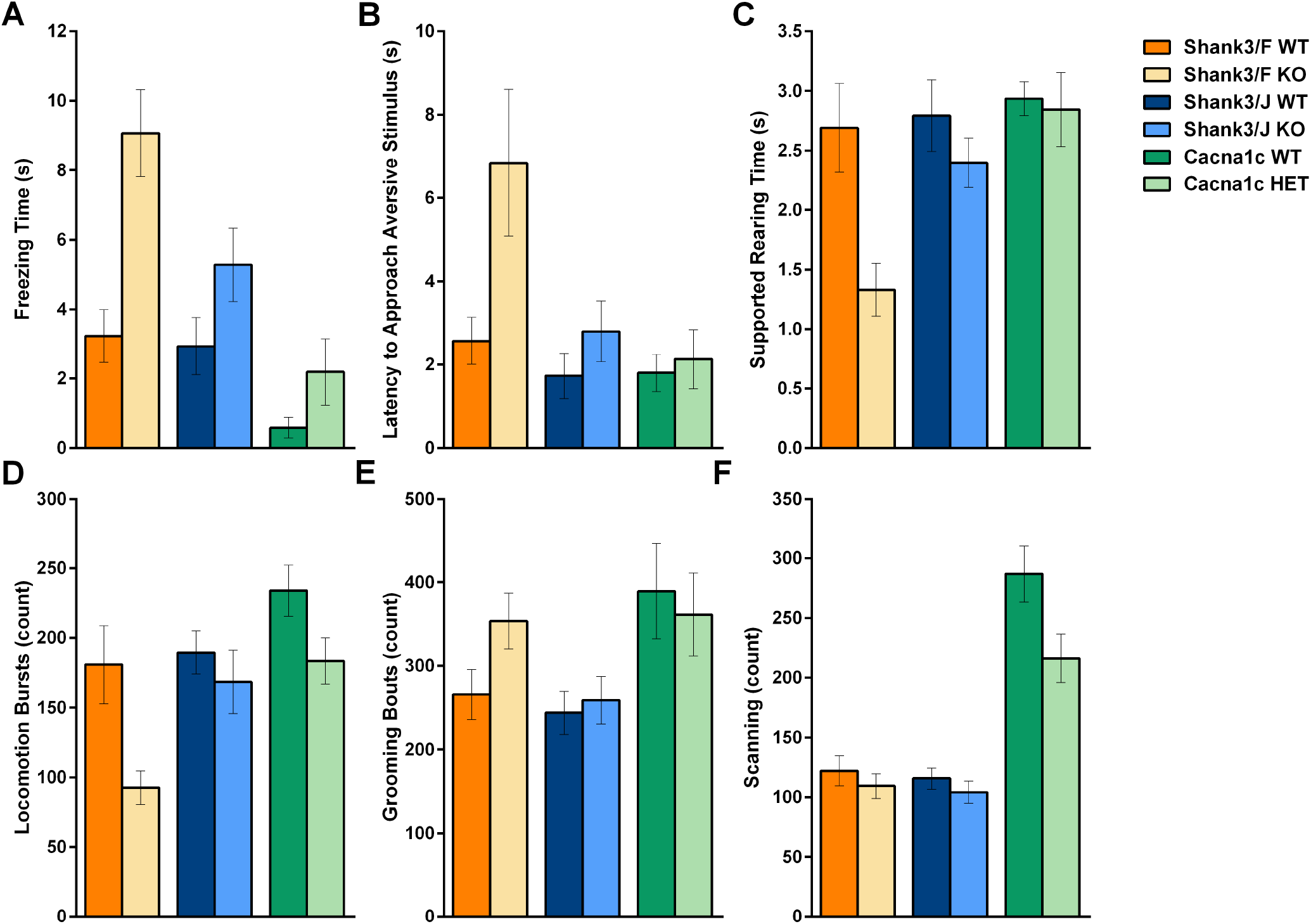
Top features in SmartCube across the three models. A: All mutant mice, Shank3/F in particular, showed some increase in freezing. B: Shank3/F KO mice showed increased latencies to approach a stimulus as compared to the WT mice. C: Shank3/F KO showed decreased rearing. D: Shank3/F KO mice, and to some extent Cacna1c HET mice, were hypoactive as compared to the WT mice. E: Shank3/F KO mice showed slightly more grooming than their WT controls. F: Cacna1c HET mice showed less scanning than WT controls. n=13-16 mice per genotype/line.

Thus, Cacna1c HET mice showed the most robust phenotype overall and *Shank3/F* KO mice showed a more robust phenotype than the *Shank3/J* KO mice, as measured by the differences from their respective WT controls. These differences were mostly driven by reduced activity and increased freezing and slightly increased grooming or sniffing.

### Neurocube

NeuroCube is a gait analysis system that automatically captures locomotion, gait geometry and dynamics, and body motion, among other features through the assessment of spontaneous locomotion in a short 5-min test session. This test has been used to probe neurological deficits in models of disease, and to assess pain and drug signatures. Although not part of the core symptomatology of autism, movement impairments have been described in a high percentage of children with autism [14, 15] along with connective dysfunction within the motor cortex [16] and it is a highly quantifiable behavior. NeuroCube features are analyzed similarly to SmartCube (see Materials and Methods; S1 Methods) but with different domains that can be analyzed separately (Table 1).

**Table 1.** NeuroCube Results at P30 and P60. Analysis of NeuroCube data showed significant differences in several domains. The Shank3/F KO mice were significantly different from the WT control mice overall, in particular for features related to paw images and position features. The Shank3/J mice were different overall at P60 and, in particular for their paw position feature. No significant differences were seen between Cacna1c HET and WT control mice overall or in key domains at P30 or P60. The numbers shown are the maximal discrimination found between the two groups. *Indicates that the associated p-value was less than .05. n=14-16 mice per genotype/line.

| Domain | Shank3/F |  | Shank3/J |  | Cacna1c |  |
| --- | --- | --- | --- | --- | --- | --- |
|  | P30 | P60 | P30 | P60 | P30 | P60 |
| **All features** | 75%* | 80%* | 56% | 79%* | 66% | 59% |
| **Gait** | 66% | 60% | 62% | 61% | 63% | 56% |
| **Imaging** | 82%* | 70% | 53% | 61% | 60% | 59% |
| **Rhythm** | 55% | 62% | 56% | 66% | 55% | 59% |
| **Body position** | 53% | 60% | 54% | 68% | 61% | 57% |
| **Paw position** | 75%* | 71%* | 57% | 88%* | 65% | 59% |

In NeuroCube, *Shank3/F* KO mice showed significant differences overall (Figs 3A and 3B; Table 1), and in particular, reductions in paw image features such as image size (at P30), and reduction in the variability of the paw position at both P30 and P60. Investigation of gait features revealed some reduction in hind base; otherwise, gait seemed to be quite normal (Fig 4; Table 1; S2 Table). Again, to provide a comparison across models, we represent the same features for the three lines, even differences were not found for all three models. It is important to note that the aggregate of many small subtle behavioral changes could contribute to an overall robust signature in NeuroCube. A spatial representation of average paw placement depicts both the slightly but consistently reduced hind base width and reduction in paw position variability (Fig 5A).

**Fig 3.**
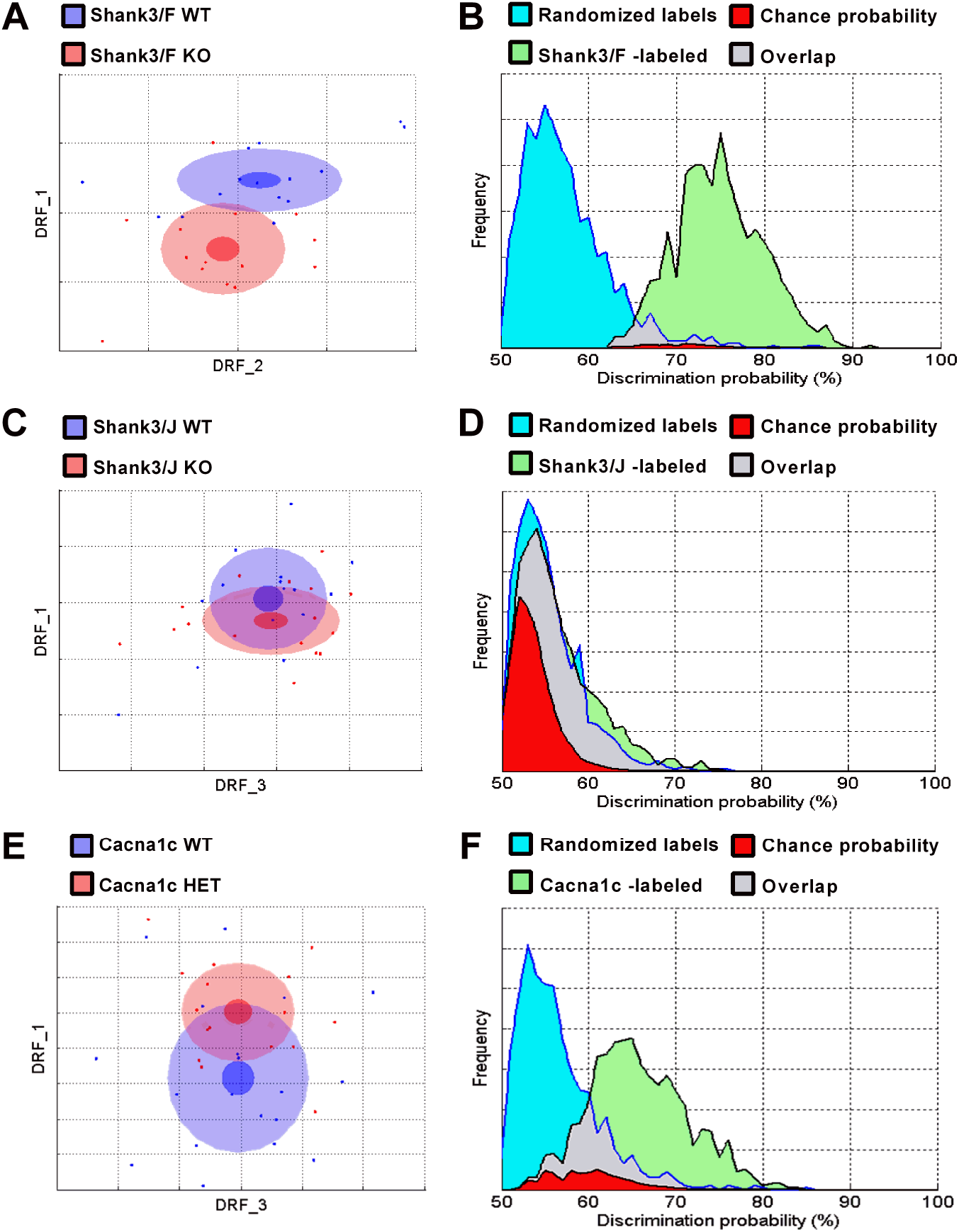
NeuroCube found different degrees of separation between mutant Shank3/F, Shank3/J, and Cacna1c mice as compared to their corresponding WT control littermates at P30 (P60 not shown). A & C: As described previously in detail [1], to build a 2D representation of the multidimensional space in which the two groups are best separated, we first find statistically independent combinations of the original features, pick the two new composite features that best discriminate between the two groups, and used them as x- and y-axes (drf 1 and 2; see S1 Methods). Each dot represents either a WT (blue) or a mutant (red) mouse. The center, small and large ellipses are the mean, standard error and standard deviation of the composite features for each group. The overlap between the groups is used to calculate the discrimination index, which measures how reliably a classifier can be trained to discriminate between the two groups (the more overlap, the worse the discrimination). B & D: To estimate how likely it is that such separation is simply due to chance, the obtained classifier is challenged many times with correctly labeled samples (green distribution) or with randomized labels (blue distribution). The overlap between these two distributions (in red) represents the probability of obtaining the observed discrimination by chance. A: At P30, the Shank3/F KO standard deviation ellipse overlapped to a small extent with the WT control ellipse. B: 75% discrimination between Shank3/F KO and WT mice could be found by chance with p < 0.02. C: At P30 the Shank3/J ellipse overlapped with the WT group almost completely, as a result of the very small phenotypic differences. D: A 56% discrimination index can be found by chance with p< 0.49. E: At P30, the Cacna1c HET standard deviation ellipse partially overlapped to with the WT control ellipse. F: 66% discrimination between Cacna1c HET and WT mice could be found by chance with p< 0.12. n=14-16 mice per genotype/line

**Fig 4.**
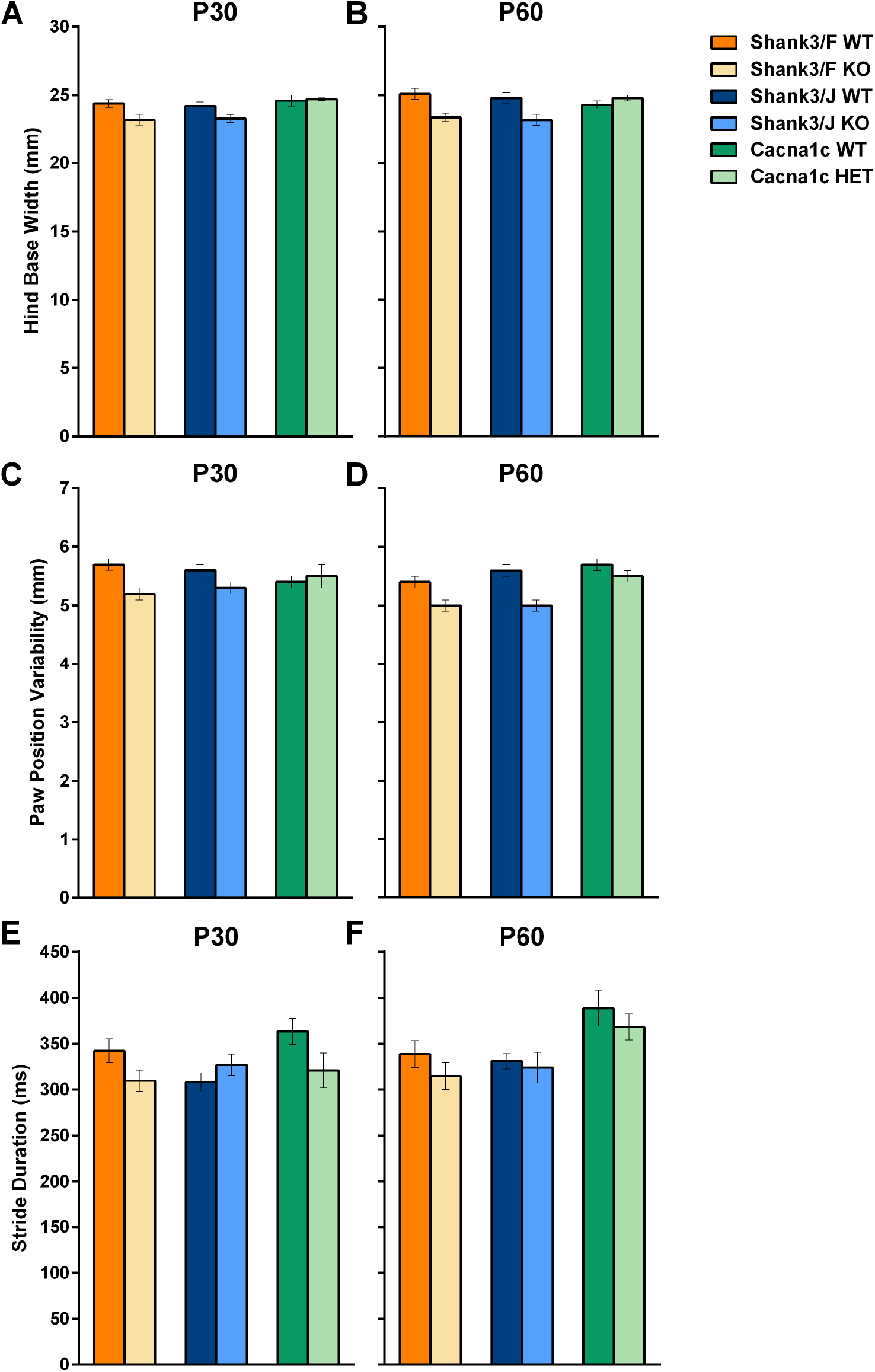
Top features in NeuroCube across the three models. A-B: The width of the base for the hind legs was reduced in both Shank3 models at the two ages studied; C-D: The variability of the position of the paw with respect to the center of the body was decreased in both Shank3 mutant models at both ages. E-F: Stride duration was reduced in the Shank3/F and Cacna1c models at P30. Data shown are means ± SEM. n=14-16 mice per genotype/line.

**Fig 5.**
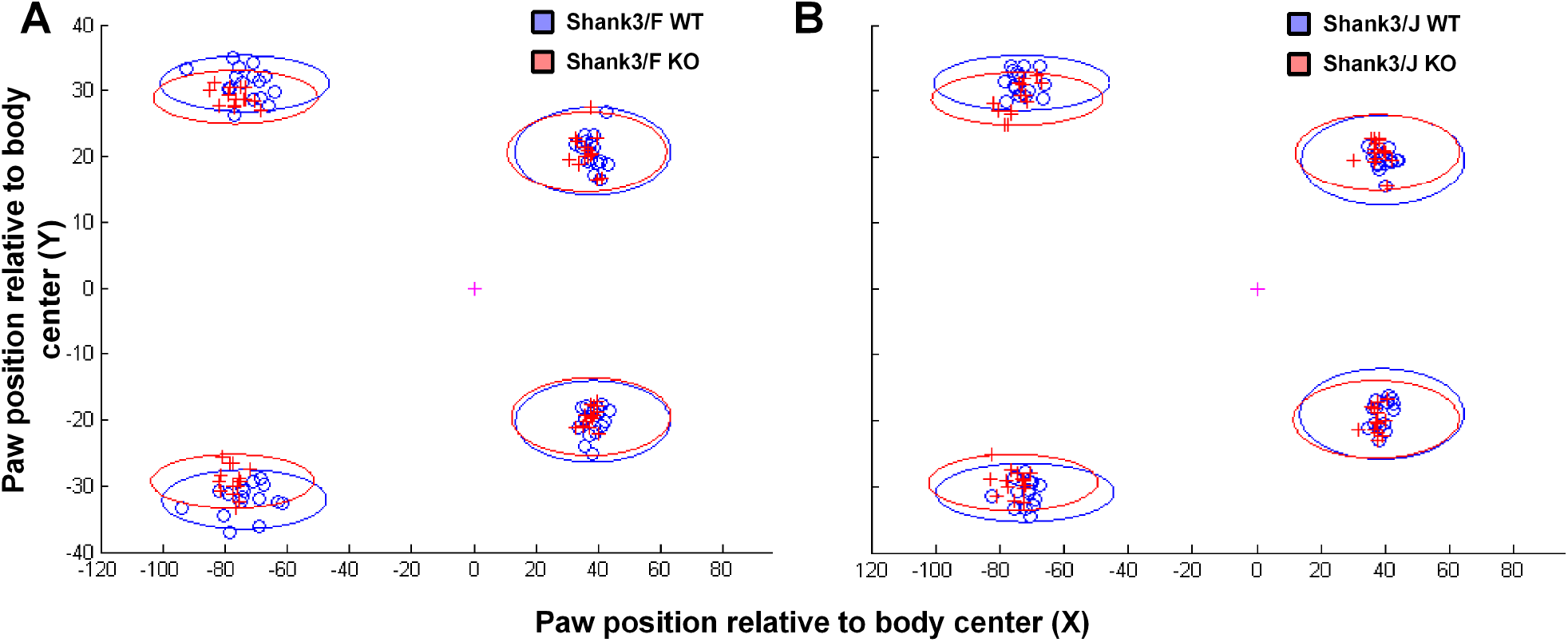
A-B: Graphical representation of the average and variability of the paw position for the four paws, for both Shank3 models at P60. The data points (plus signs and open circles) represent the individual X-Y coordinates corresponding to the paw positions of the hind limbs (on the left) and forelimbs (on the right). Ellipses show the standard deviation of the X-Y coordinates (with the long and short axes being parallel and orthogonal to the direction of movement, respectively). Ellipses corresponding to the mutant hindlimb paws mice lie on the inside of those corresponding to the WT mice, showing narrowing of the hind base. The cross in the center of the graphs represent the body center as determined by the computer vision algorithm. n=14-16 mice per genotype/line.

*Shank3/J* mice did not show robust differences overall at P30 (Figs 3C and 3D; Table 1), but showed consistent differences in paw position at P60 (Fig 4; Table 1). Further investigation of top individual features, however, revealed a somewhat reduced hind base and paw position variability, not unlike the *Shank3/F* model (Fig 4; Table 1; S2 Table). As the differences are small at P30, the machine learning algorithm failed to separate the groups, although the (weak) pattern seems evident *a posteriori*. Fig 5B depicts spatially the reduced hind base.

Cacna1c HET mice did not show significant differences compared to WT mice overall at P30 (Figs 3E and 3F) or P60 (Table 1). Moreover, key domain areas of gait, paw imaging, rhythm, body position, and paw position also seemed to be quite normal (Table 1; S2 Table). Stride duration was slightly reduced in Cacna1c HET mice at P30 compared to WT mice, but few other differences were observed in this test (Fig 4; S2 Table).

As variability in speed affects gait measures [17], we investigated the correlations between speed and top gait features at P60 (Figs 6A and 6B). The top features were somewhat different for the *Shank3* models than for the Cacna1c model, leading to the different analyses depicted in Fig 6. In the *Shank3* models, not only were correlations within groups weak (with slopes not significantly different from zero; *p’*s > .05), but also WT and KO regression lines were parallel to each other (see Figs 6A and 6B), showing that genotypic differences existed over and above minor correlations with speed (only the regression for paw print variability versus speed reached significance for the *Shank3/J* KO mice; *p*< 0.02). Top features found to contribute more robustly to the discrimination between groups were graphed to interpret the bioinformatics’ results but not further analyzed statistically.

**Fig 6.**
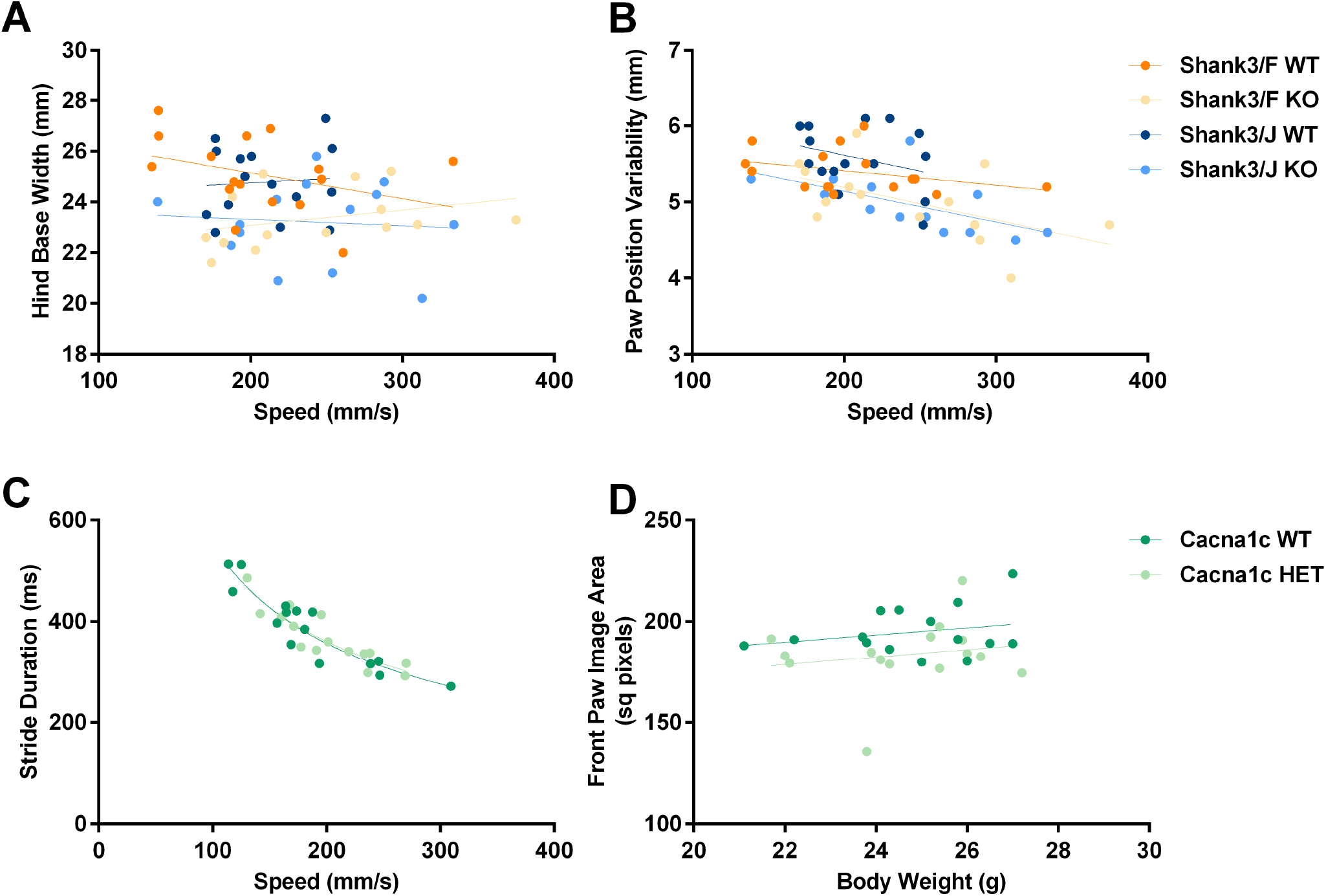
Correlation analysis of gait features as a function of speed or body weight. A: At P60 the width of the hind base as a function of speed did not show consistent correlations. Both Shank3 WT groups were above the mutant groups suggesting that their narrower base was not an artifact of a somewhat faster speed; B: The variability of the paw position seemed to decrease as a function of speed; the data corresponding to the two Shank3 WT groups and the fitted regression lines, however, were above the two mutant groups, again suggesting that the decreased paw position variability was decreased in the mutants over and above the effect of speed. C-D: Stride duration was non-linearly correlated with locomotor speed. A power regression provided the best fit to these dependencies and suggested a log-log transformation for statistical analysis of differences between regression lines to assess to the extent to which changes in gait dynamics could be secondary to increased locomotor speed. The front paw image was linearly related to body weight, suggesting that this measure was affected by body weight rather than indicative of neurological dysfunction. n=14-16 mice per genotype/line.

In the Cacna1c model, no differences in stride duration were observed between the WT and mutant mouse data after log-log transformations were performed to ensure linearity (Fig 6C). Thus, it appears that a reduction in stride duration was affected by faster speed within both groups. As described previously, differences in paw imaging may be caused by neurological deficit or secondary to a reduced body weight [1]. The front paw image area, visualized as a scattergram as a function of body weight, shows that Cacna1c HET mice did not differ from WT mice in size, nor were there differences in paw image area (Fig 6D). Although robust regressions were observed within each group, differences between the regression lines of the two groups were not significant, indicating no genotypic differences.

Thus, both *Shank3* models seem to have a narrower base and reduced variability. These characteristics do not seem to resemble any disease-typical gait, which generally comprises the opposite features, namely, wider base and higher variability. The Cacna1c HET mice appear to be similar to WT mice in this test.

## Development

Deviations in general health and in the development of rudimentary behaviors in early life may be early indicators of later pathology. In the first week of life, simple motor behaviors emerge, physiological measures such as thermoregulation and body weight may show genotypic differences, and anxiety-like behaviors (i.e. USVs and pivoting) are expressed to varying degrees [18]. Increasingly independent behaviors such as grooming appear around P9 and, after two weeks of age, physiological and anxiety-like measures correlate as the adrenal-pituitary axis matures [19].

### Body weight

All groups gained weight with age during both the neonatal and later periods (Fig 7; S3 Table). *Shank3/F*, but not *Shank3/J* or Cacna1c, mutant mice showed slight but significantly higher weight beginning at P60, as compared to their control mice.

**Fig 7.**
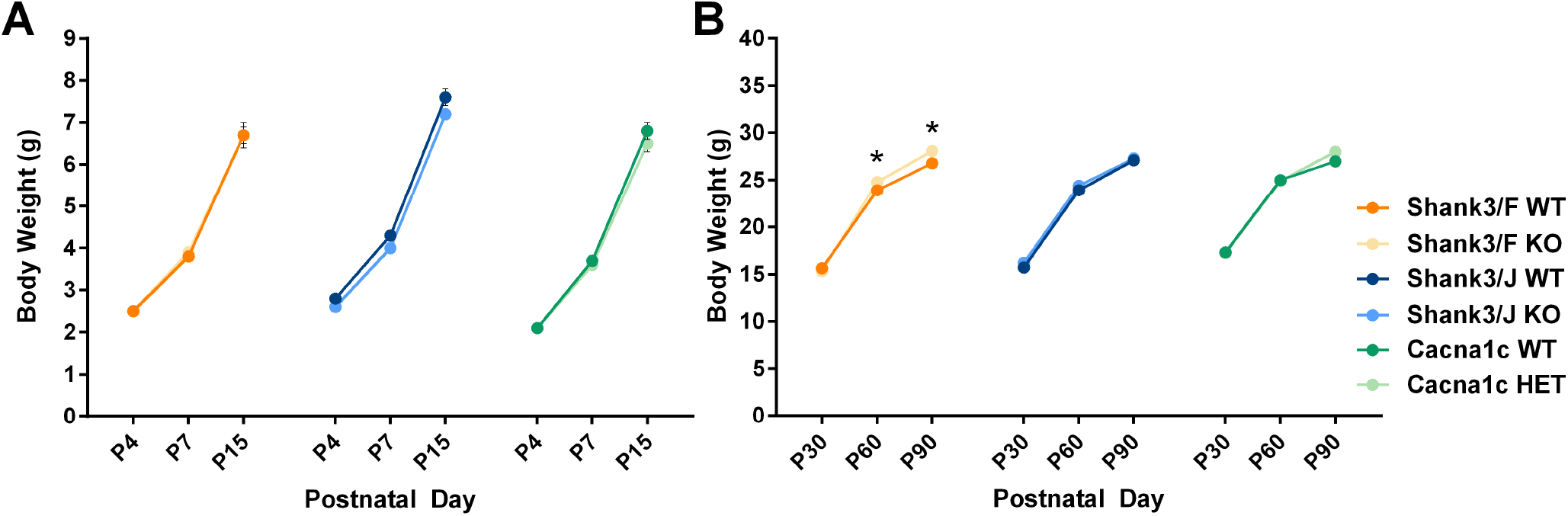
Body weight during the neonatal (A) and postnatal (B) periods. Simple main effect analysis following a significant Genotype X Age interaction showed significant differences for the Shank3/F model after P60 as compared to corresponding WT controls. Note errors bars are hidden by the graphing symbols. Data shown are means ± SEM; n=15-16 mice per genotype/line. (*p< 0.05)

### Milk content

During the first post-natal days this is a sign that pups are able to nurse and therefore an indicator of early health. Stomach milk was apparent at P4 and P7 in most *Shank3* and in all Cacna1c pups with no genotypic differences seen at either age (S3 Table).

### Eye opening

About half of the pups in the *Shank3* groups had eyes open at P13 and no genotypic differences were seen (S3 Table). Few pups of either genotype had both eyes open at P13 in the Cacna1c model, with fewer mutant pups with both eyes open than WT pups (S3 Table).

### Ultrasonic vocalizations (USVs)

USVs serve a functional purpose as they trigger a dam’s retrieval response when a pup is outside the nest [20-22]. Call rates in the *Shank3* models were very low at all ages, quite minimal at P4 for all models, and then increased up to P15, particularly in the Cacna1c model where they showed a significant increase with age (Fig 8A). No genotypic differences were found (S4 Table).

**Fig 8.**
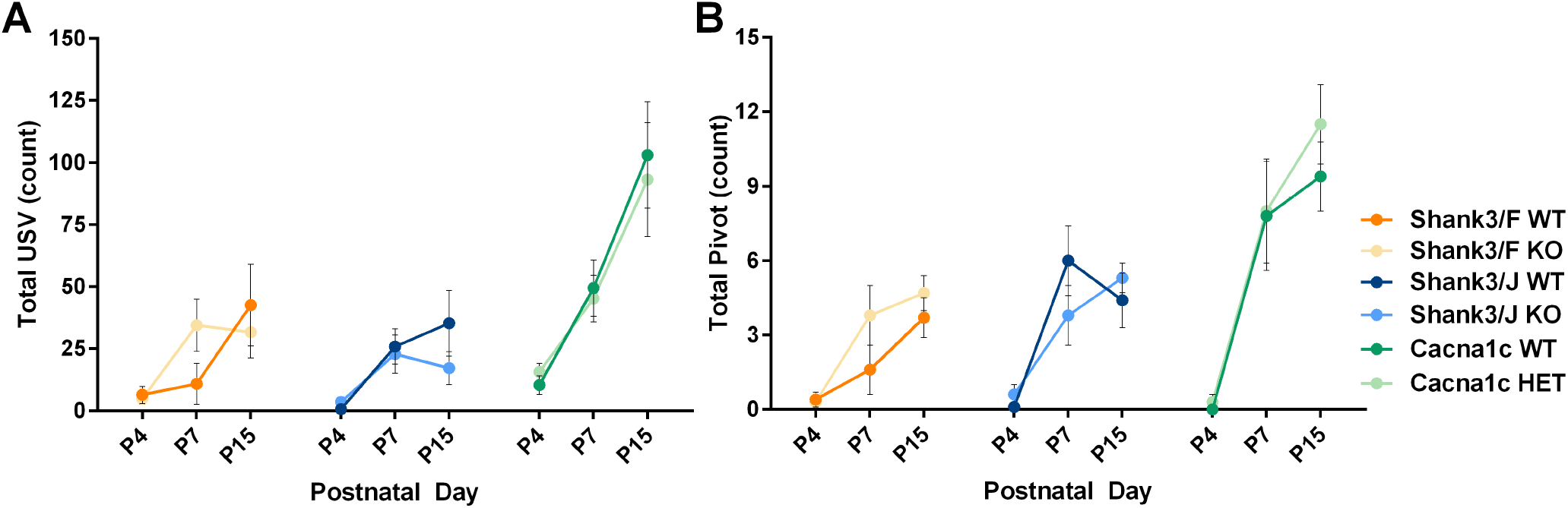
During the isolation test mice were separated from the dams and individually tested in a clean cage. A: There were no genotypic differences in neonatal ultrasonic vocalizations (USVs) for any of the three models. B: Pivoting, normally associated with the production of USVs, did not show genotypic differences either. Data shown are means ± SEM; n=15-16 per genotype/line.

### Activity

In both lines, locomotor activity, as measured by square crossings, significantly increased with age but no genotypic differences were observed (S1A Fig; for all data see S4 Table). Pivoting is associated with the number of ultrasonic calls and thought to help to propagate the sound in a wider range and interestingly we found these two measures showed similar patterns (Fig 8B). Rearing was very rare until P15, when pups have better motor coordination. At P15, only the *Shank3/J* KO mice showed increased rearing. Olfaction is a primary sensory modality in mice and sniffing was observed at all ages with no genotypic differences. Grooming was infrequent and present only at P15 with no genotypic differences seen.

Motor difficulty seems to be an early sign in autism and related neurodevelopmental disorders [23]; therefore it is relevant to measure motor development in animal models. Overall, motor coordination appeared normal in all groups though, at P15, Cacna1c HET mice walked down less than WT mice but turned more completely in the geotaxis test (Table 2; S5 Table). There were no genotypic differences in righting reflex measures, a response to changes in proprioception, and rolling onto one side was very rarely observed for any of the three models (Fig 9). The geotaxis test allows measuring motor coordination and, during the first week of development, the turning response to gravitational pull. During normal development the early turning response disappears and other behaviors, such as walking sideways or downwards, or jumping, appear. Other observed yet infrequent behaviors included stretch-attend, jump, and twitch. No apparent differences were present for these rare behaviors.

**Fig 9.**
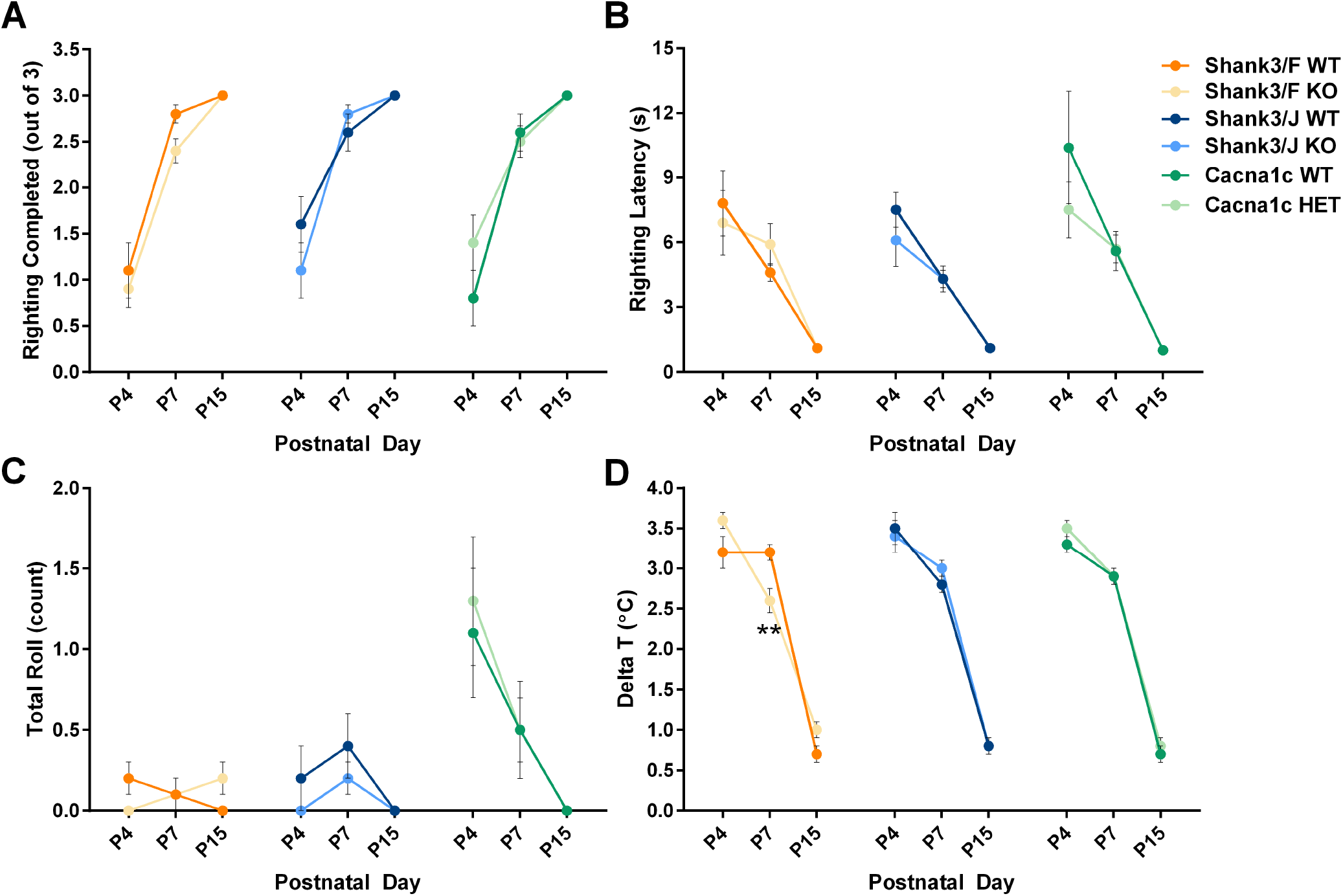
During the neonatal test, a number of motor coordination measures are taken in addition to thermoregulation. In the righting test, mice are placed on a surface on their backs and must maneuver to a position where they are on all four limbs. The frequency of falling over or rollling is also used as a measure of motor coordination in the isolation test. Thermoregulation is assessed by a measure of axillary temperature taken before and after the isolation test. A: Number of righting successes. B: Righting latency. C: Number of rolls on a side. D: Difference between the axillary temperature measurements before and after the isolation test showed a decrease in the Shank3/F temperature at P7. Data shown are means ± SEM; n=15-16 mice per genotype/line. (**p < 0.01)

**Table 2.** Various outcome measures of the geotaxis test across development. Mice were placed on an inclined plane and their behavior was observed. A fall indicates lack of coordination whereas a turn indicates a geotactic reflex response. After P7 this reflex is lost and other behaviors appear, such as walking down the plane. *Indicates a significant genotypic difference in the highlighted age/measure. n=15-16 mice per genotype/line.

|  | Shank3/F WT |  |  | Shank3/F KO |  |  | Shank3/J WT |  |  | Shank3/J KO |  |  | Cacna1c WT |  |  | Cacna1c HET |  |  |
| --- | --- | --- | --- | --- | --- | --- | --- | --- | --- | --- | --- | --- | --- | --- | --- | --- | --- | --- |
| Age | P4 | P7 | P15 | P4 | P7 | P15 | P4 | P7 | P15 | P4 | P7 | P15 | P4 | P7 | P15 | P4 | P7 | P15 |
| % falls | 53.1 | 34.4 | 0.0 | 56.3 | 31.3 | 0.0 | 40.6 | 21.9 | 0.0 | 43.8 | 15.6 | 0.0 | 25.0 | 6.3 | 0.0 | 36.7 | 10.0 | 0.0 |
| % turns | 18.8 | 31.3 | 62.5 | 0.0 | 28.1 | 87.5 | 0.0 | 21.9 | 43.8 | 6.3 | 56.3 | 71.9 | 0.0 | 6.3 | 50.0* | 3.3 | 26.7 | 100.0* |
| % walk down | 0.0 | 0.0 | 25.0 | 0.0 | 0.0 | 12.5 | 0.0 | 3.1 | 34.4 | 0.0 | 0.0 | 28.1 | 0.0 | 0.0 | 37.5* | 0.0 | 0.0 | 0.0* |
| % no reaction | 28.1 | 34.4 | 12.5 | 43.8 | 40.6 | 0.0 | 59.4 | 53.1 | 21.9 | 50.0 | 28.1 | 0.0 | 75.0 | 87.5 | 12.5 | 60.0 | 63.3 | 0.0 |

### Thermoregulation

Homeostatic regulation is weak during the first postnatal week [19]. Pups lose temperature when separated from the litter during the 3-4 minute handling and observational test. The *Shank3/F* KO mice lost slightly less heat than their corresponding WT mice at P7 but were otherwise similar to WT mice at other time points (Fig 9D; S1B Fig; S3 Table). There were no significant differences between genotypes for the *Shank3/J* or Cacna1c mouse models (Fig 9D; S1B Fig; S3 Table).

## Tests of Social Behavior

### Three-chamber test

The three-chamber test is one of the most widely used behavioral tests to assess social behavior [24]. During the habituation phase, the *Shank3/F* model, the KO mice showed less exploration of the two side chambers over the center area, as compared to the WT mice (S2 Fig; see S6 Table for all statistics). There were no differences between genotypes for the *Shank3/J* or Cacna1c mice, but both the WT and mutant mice of the Cacna1c model explored one side of the chamber more than the other.

During the social preference phase, *Shank3/F* WT mice seemed to prefer, as expected, to spend more time in the social than in the object chamber whereas KO mice did not show any differential exploration. This apparent genotypic effect, however, did not reach significance as the Genotype × Chamber side interaction yielded a p-value of 0.29 (Figs 10A, 10B and 11A; S2 Fig). However, the *Shank3/F* WT mice did not show as great a preference as did the *Shank3/J* WT mice, contributing to this failure to reach significance. Both *Shank3/F* WT and KO mice sniffed the mouse longer than the object similarly suggesting that if a social deficit exists in *Shank3/F* KO mice, this characteristic is rather subtle. We did not observe sociability deficits in the *Shank3/J* or Cacna1c models; both genotypes preferred, to an equivalent extent, to spend more time in the social chamber than in the object chamber, and to sniff the mouse longer than the object. Activity, as indicated by the entries into the side chambers did not differ across stimuli or genotypes in the *Shank3/F* or Cacna1c models, but *SHANK3/J* mice and their control littermates crossed over to the mouse chamber more than to the object chamber (S2 Fig).

**Fig 10.**
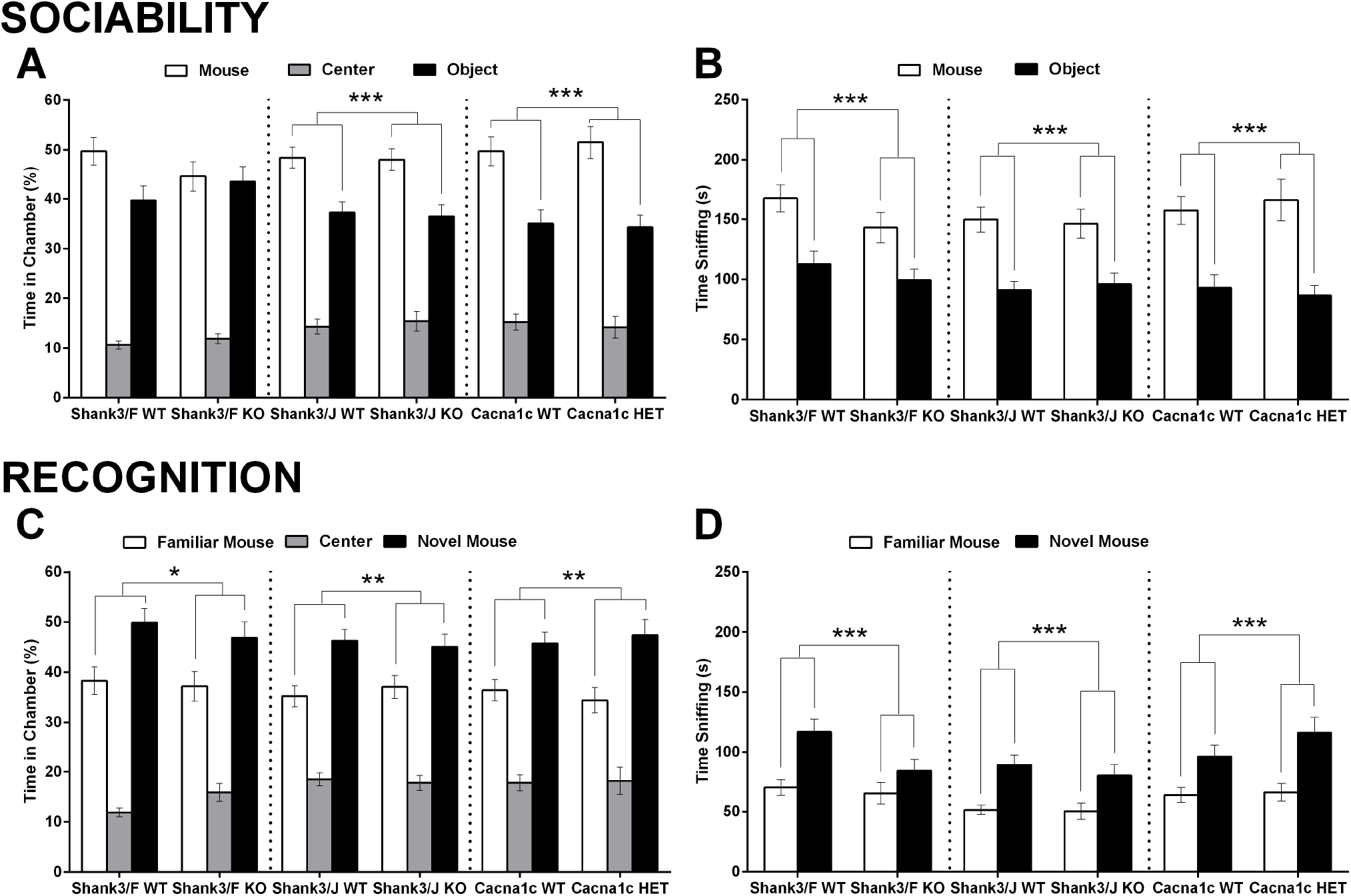
The three-chamber test assesses preference for a social versus a non-social stimulus (“Sociability”) and recognition of a novel versus a familiar social stimulus (“Recognition”). A. During the sociability phase, Shank3/F KO mice did not show a preference for the social stimuli. The difference with the WT group, however, did not reach significance. Shank3/J and Cacna1c mutant and WT groups spent more time in the social than the object chamber to the same extent, suggesting intact social preference. B. During the sociability phase, all groups sniffed the social container more than the object container to a similar extent. C. During the recognition phase, all groups spent more time in the novel than in the familiar mouse chamber, again to a similar extent. D. During the recognition phase, all groups sniffed the novel more than the familiar social container to a similar extent. Time in each chamber was recorded automatically, whereas sniffing was scored manually. Data shown are means ± SEM; n=16 mice per genotype/line. (Chamber side main effect: *p< 0.05, **p< 0.01; ***p< 0.001.)

**Fig 11.**
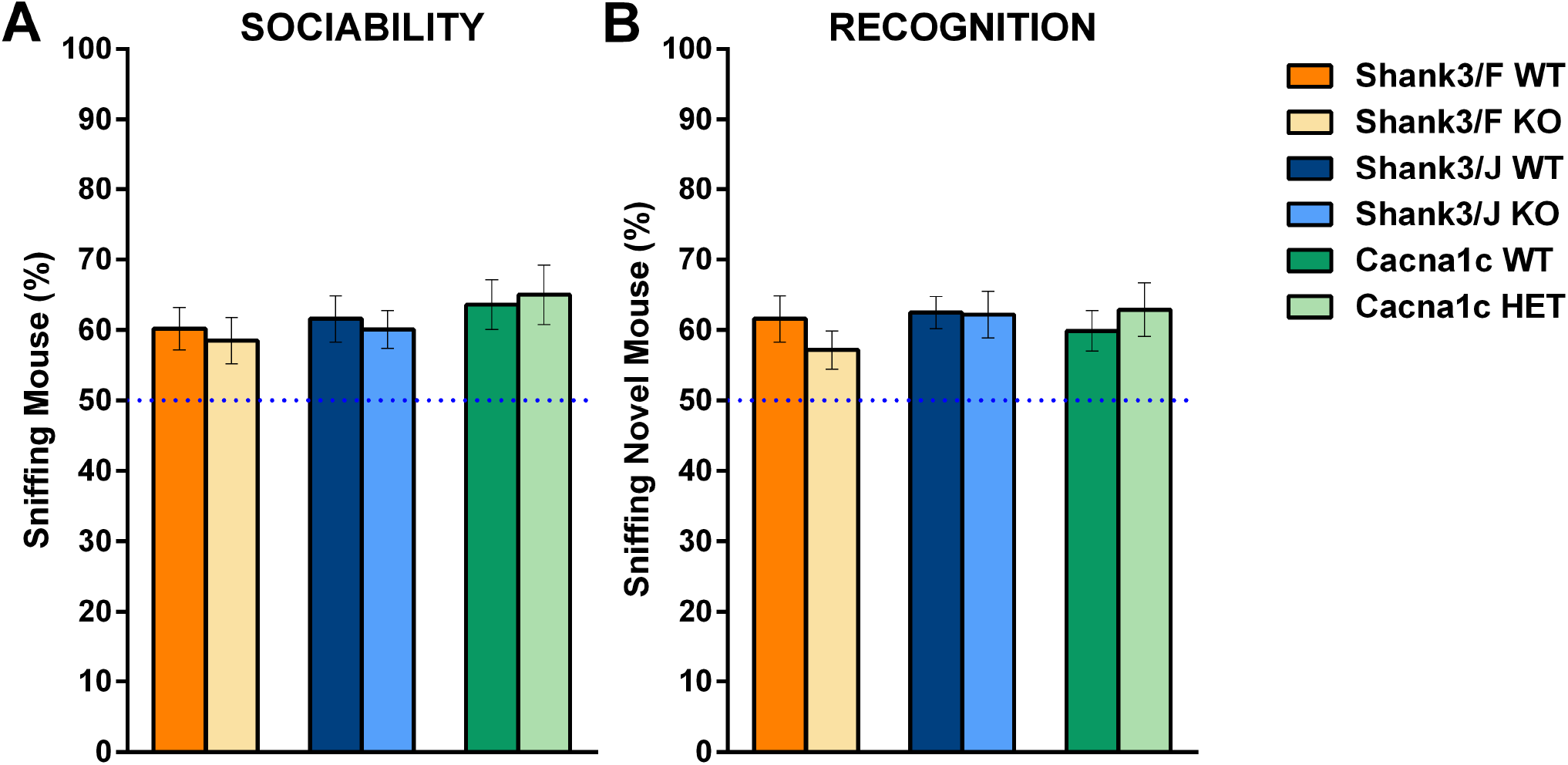
In the three-chamber test, preference was calculated by the proportion of the total time sniffing spent with the mouse or with novel stimuli, with a dashed line at 50% indicating a lack of preference. A: All groups showed preference for sniffing the mouse over the object. B: All groups showed preference for sniffing the novel mouse over the familiar mouse. Sniffing was scored manually. Data shown are means ± SEM; n=16 mice per genotype/line.

Overall, none of the models showed deficits in social recognition. All groups preferred, to an equivalent extent, to spend more time in the novel mouse chamber than in the familiar mouse chamber, and to sniff the novel mouse longer than the familiar mouse (Figs 10C, 10D and 11B; S2 Fig). Although *Shank3/F* KO mice showed fewer entries into the two side chambers than their control mice, entries into the novel mouse chamber was higher than into the familiar mouse chamber for all groups to the same extent. Chamber crossings were not significantly different between the *Shank3/J* mutant and WT mice and were higher to the chamber containing the novel mouse. Cacna1c mutant mice showed fewer entries into the two side chambers than their control mice and neither genotype showed a difference in chamber crossings (S2 Fig).

*Shank3/F* KO mice, therefore, show subtle social deficits, with only one of several social measures suggestive of a deficit. *Shank3/J* and Cacna1c mutant mice, on the other hand, show normal social preference on all measured behaviors in this test.

### Reciprocal social interaction

In the reciprocal social interaction assay, pairs of genotype- and age-matched mice are allowed to freely interact. *Shank3/F* mutant pairs spent more time in close proximity with each other and engaged in reciprocal interactions for more time than the corresponding WT pairs (Fig 12A; S4 Fig). Overall, *Shank3/J* mutant mice showed normal social interactions. Their time in close proximity was similar to that shown by the WT control pairs (Fig 12A). The only significant different type of interaction was that *Shank3/J* KO mice engaged in nose-center contact for more time compared to WT pairs (Fig 12C). Cacna1c mutant pairs spent more time in close proximity with each other and engaged in nose-back contact for more time compared to WT pairs (Figs 12A and 12C). Consistent with the hypoactive profile obtained in SmartCube, all mutant groups moved less and spent significantly less time following their partner than their corresponding WT groups (S3 Fig; see all statistical results in S7Table). However, the distance between the paired subjects was similar among all groups (Fig 12B). There were no genotypic differences in the rates of active, passive and reciprocal interactions (Fig 12D).

**Fig 12.**
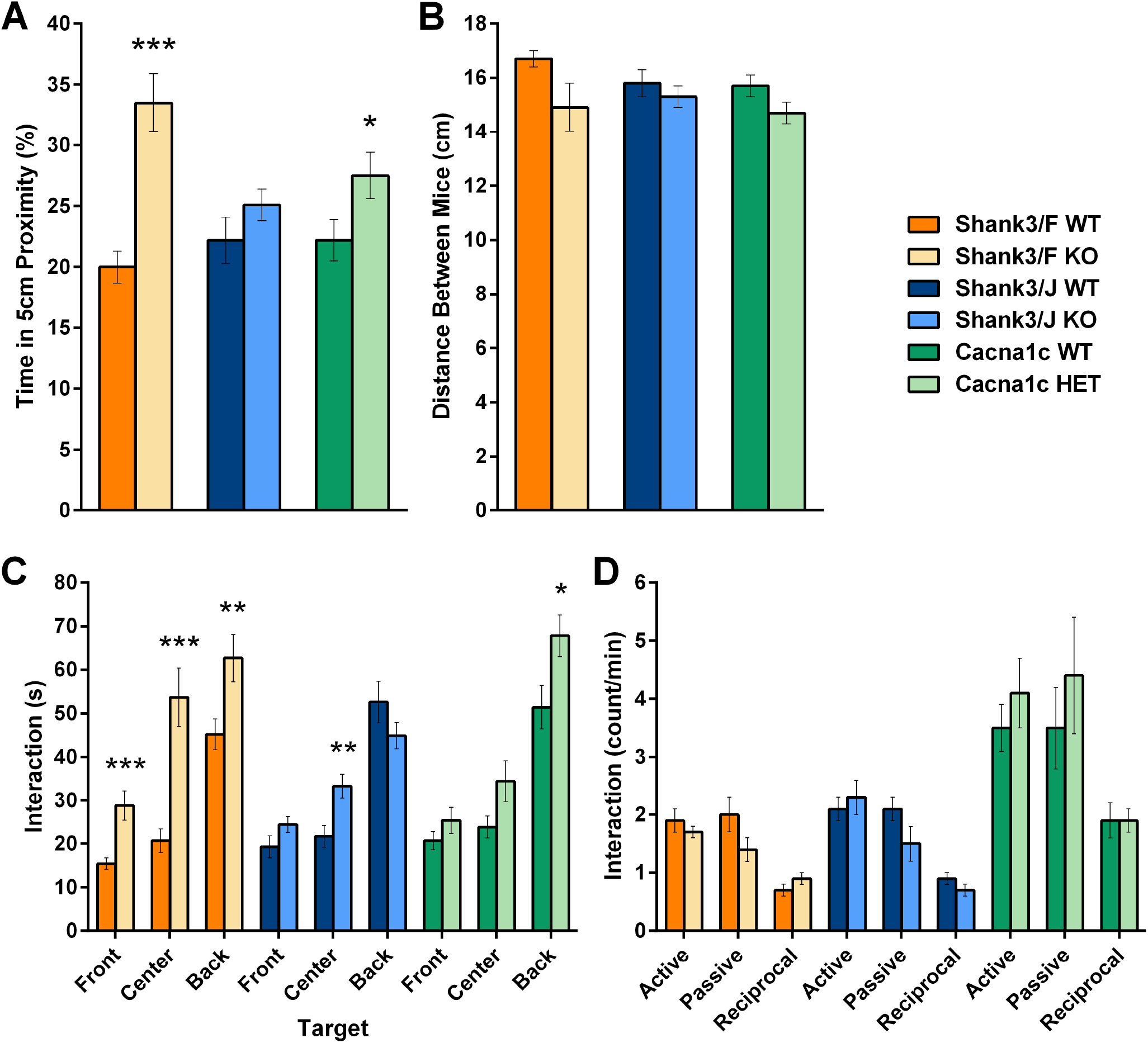
The reciprocal social interaction test did not show social deficits in any of the models. A: The time in which the mice were closer than 5 cm was increased in the Shank3/F and Cacna1c but not the Shank3/J mutant pairs as compared to their respective WT controls. B. Mutant mice pairs did not show a difference compared to their corresponding WT control pairs in the average distance maintained between the pairs of mice. C: Duration of nose-to-front, -side or –back contacts with the paired mouse. Shank3/F mutant mice showed increased interaction towards their partner’s front, center, and back. Shank3/J mutant mice increased in the amount of time interacting to the center compared to WT mice. Cacna1c mutant mice interacted to their partners’ back more than WT mice. D: Active, passive and reciprocal interactions were similar between mutants and their respective controls. All behaviors with the exception of the active, passive and reciprocal interactions were recorded using an automated system. Data shown are means ± SEM; n=16 mice per genotype/line. (*p< 0.05, **p< 0.01, ***p< 0.001)

Vocalizations were not very frequent in these testing conditions and showed no genotypic difference. In general, therefore, the three models appeared hypoactive with increased duration of some social behaviors but decreased active pursuit of interactions.

### Urine-exposure open field

Male mice mark territories and vocalize in response to the scent of urine from an estrous female. Scent-marking requires no social experience [25, 26], whereas ultrasonic vocalizations require social experience with females and female scent exposure. There were no differences between the *Shank3/F* KO and WT control mice in total locomotion and locomotion in the center during the baseline session (see all statistics in S8 Table), but there was a significant reduction in distance traveled (overall and in the center) during the exposure session. *Shank3/F* KO mice spent less time in the center during both the baseline and exposure sessions (Figs 13A and 13B), suggesting an anxiety-like phenotype, and vocalized less than the WT control mice (Fig 13C). There were no differences in the number and amount of scent marks between *Shank3/F* KO and WT mice, though scent-marking area was smaller during the exposure session (Fig 13D). *Shank3/J* mice also showed no differences in total locomotion and locomotion in the center during the baseline and exposure sessions (see all statistics in S8 Table). However, *Shank3/J* KO mice spent less time in the center during both sessions (Figs 13A and 13B) and vocalized less than the WT control mice (Fig 13C). The number and amount of scent marks were similar between the *Shank3/J* KO and their corresponding WT mice but scent marking area was smaller during the second session (Fig 13D). Cacna1c HET mice traveled less distance compared to WT control mice during the baseline and exposure sessions both overall and in the center of the chamber (see all statistics in S8 Table). Cacna1c HET mutant mice spent less time in the center during the exposure session and tended to spend less time in the center during the baseline session (Figs 13A and 13B). Although the number of ultrasonic vocalizations seemed reduced compared to WT control mice (Fig 13C), the difference did not reach significance. There were no differences in the number and size of scent marks between Cacna1c HET and WT mice, though scent-marking areas were smaller during the exposure session (Fig 13D).

**Fig 13.**
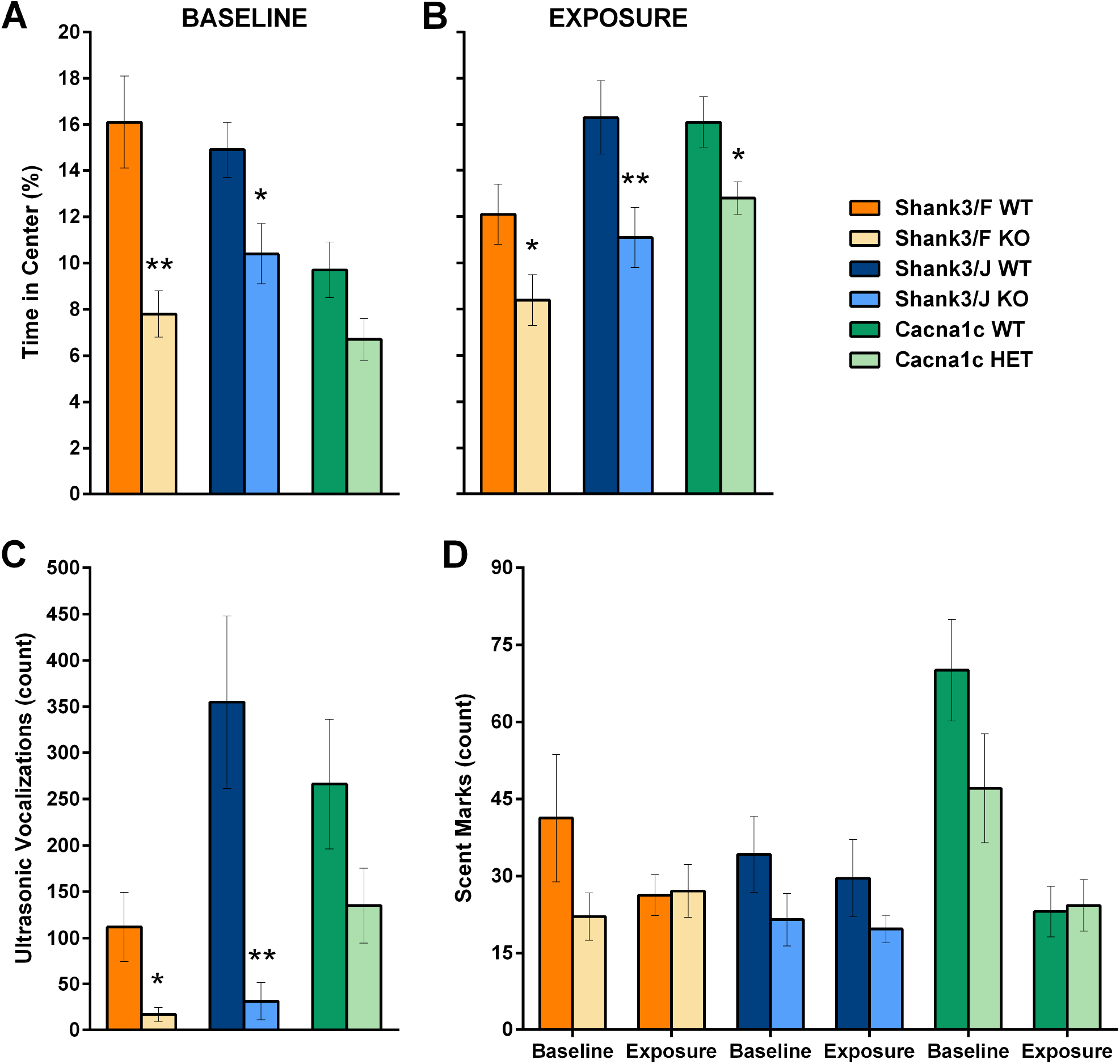
In the urine-exposure open field, male mice are habituated to an arena and before a small quantity of estrous female urine is placed in the center of the arena. A-B: Shank3 mutant mice explored the center of the arena less than the WT mice during (A) baseline and all models explored the center arena less during the (B) exposure session; C: Ultrasonic vocalizations emitted during the exposure session showed deficits in both mutant Shank3 models as compared to their WT controls; D: No differences were found in scent marking during baseline or urine exposure. Data shown are means ± SEM; n=16 mice per genotype/line. (*p< 0.05; **p< 0.01)

All models therefore show an anxiety-like phenotype, with the *Shank3* models showing a clear reduced social response to a female stimulus.

## Cognition

### Procedural T-maze

The procedural T-maze is an egocentric visuomotor task, variations of which have been used to understand cortico-striatal and hippocampal circuits [27, 28]. The reversal phase assesses behavioral flexibility, as animals need to inhibit a preponderant response to accurately perform under a new rule. There were no significant effects due to genotype for any of the models (Fig 14A; see S9 Table). During reversal, all groups reached criteria at the same rate and showed similar performance accuracy, although Cacna1c HET mice seemed to perform slightly worse during reversal (Fig 14B; S5 Fig).

**Fig 14.**
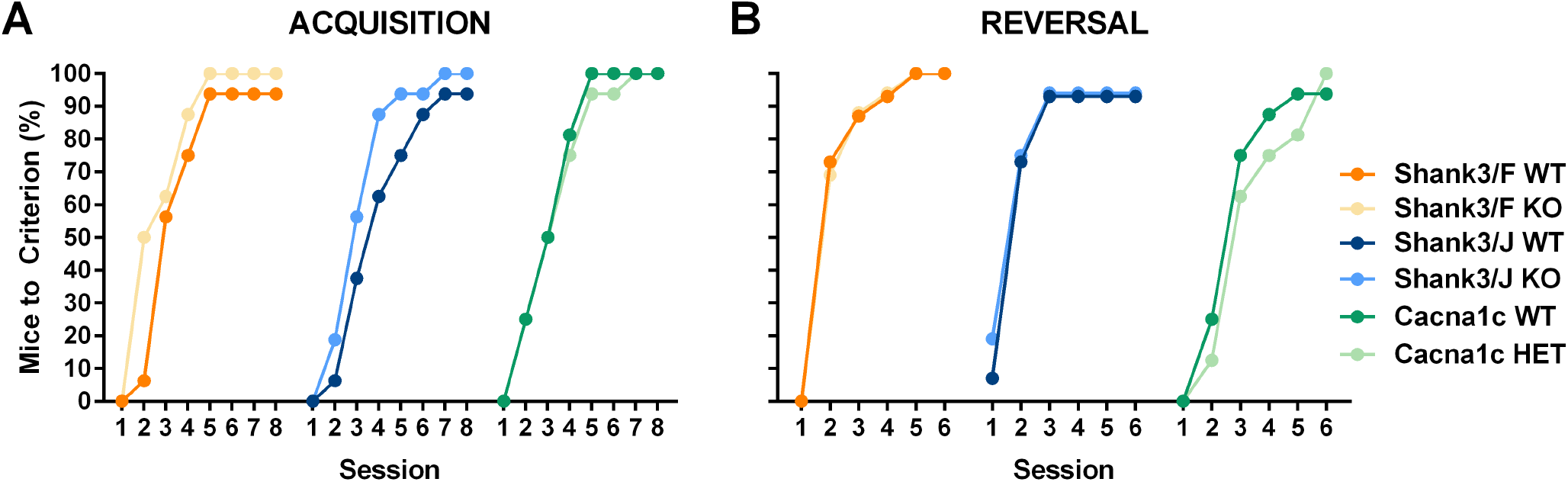
Acquisition and reversal in the procedural T-maze. Mice are first trained to reach a platform on one side of the maze. Once mice achieve 6 correct out 8 trials in two consecutive days, the platform position is reversed and mice are trained under the new rule. A: The proportion of mice that reached criteria during the acquisition (maximum 8 days) shows that there were no deficits in any of the models. B: During reversal (6 days maximum), there were no deficits either. Note the reversal data for WT mice are exactly under the Shank3 mutant mice data. Data shown are proportions of mice that started the task. n=16 mice per genotype/line.

## Sensory-Motor Gating

### Prepulse inhibition of startle

Acoustic startle is the unconditioned response to a loud stimulus. This response can be attenuated if preceded by a quieter stimulus, thus termed prepulse inhibition of startle and is considered a test of pre-attentive processing [29-31]. *Shank3/F* KO mice showed significantly reduced startle and enhanced prepulse inhibition compared to their WT controls, whereas *Shank3/J* and Cacna1 mutant mice and WT controls were similar in both measures (Fig 15; S10 Table).

**Fig 15.**
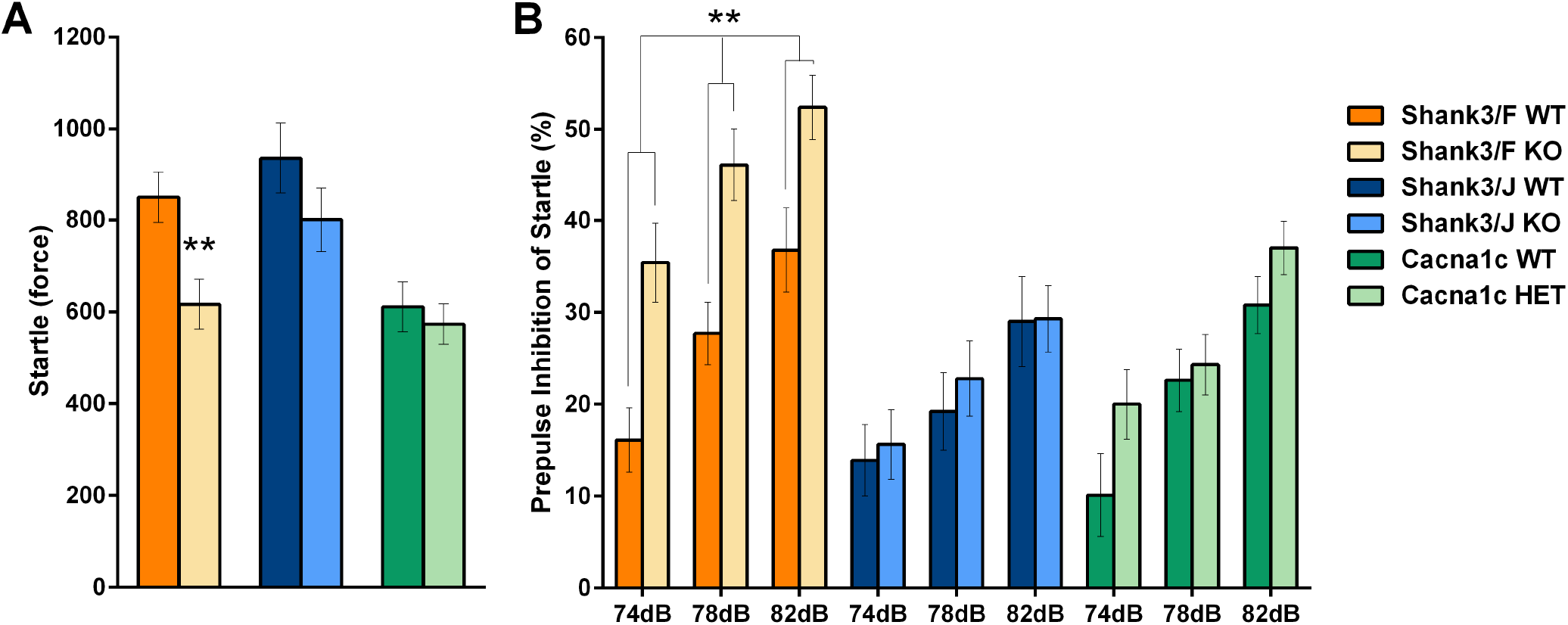
Response to startle and prepulse inhibition of startle. A: Shank3/F but not Shank3/J or Cacna1c mutant mice showed decreased startle responses (force arbitrary unit) compared to their WT control mice. B: Shank3/F but not Shank3/J or Cacna1c mutant mice showed higher prepulse inhibition of startle than the corresponding WT control mice. Data shown are means ± SEM; n=16 mice per genotype/line. (**p< 0.01)

## Repetitive/Anxiety-like Behavior

### Marble burying

Behavioral responses to novel diggable media can be used to assess anxiety-like, stereotypic and/or obsessive-compulsive-like behavior in models of autism [32-34]. *Shank3/F* KO mice buried significantly fewer marbles and showed reduced locomotor activity whereas *Shank3/J* and Cacna1c mutant mice were similar to their WT controls in both measures (Fig 16; S11 Table).

**Fig 16.**
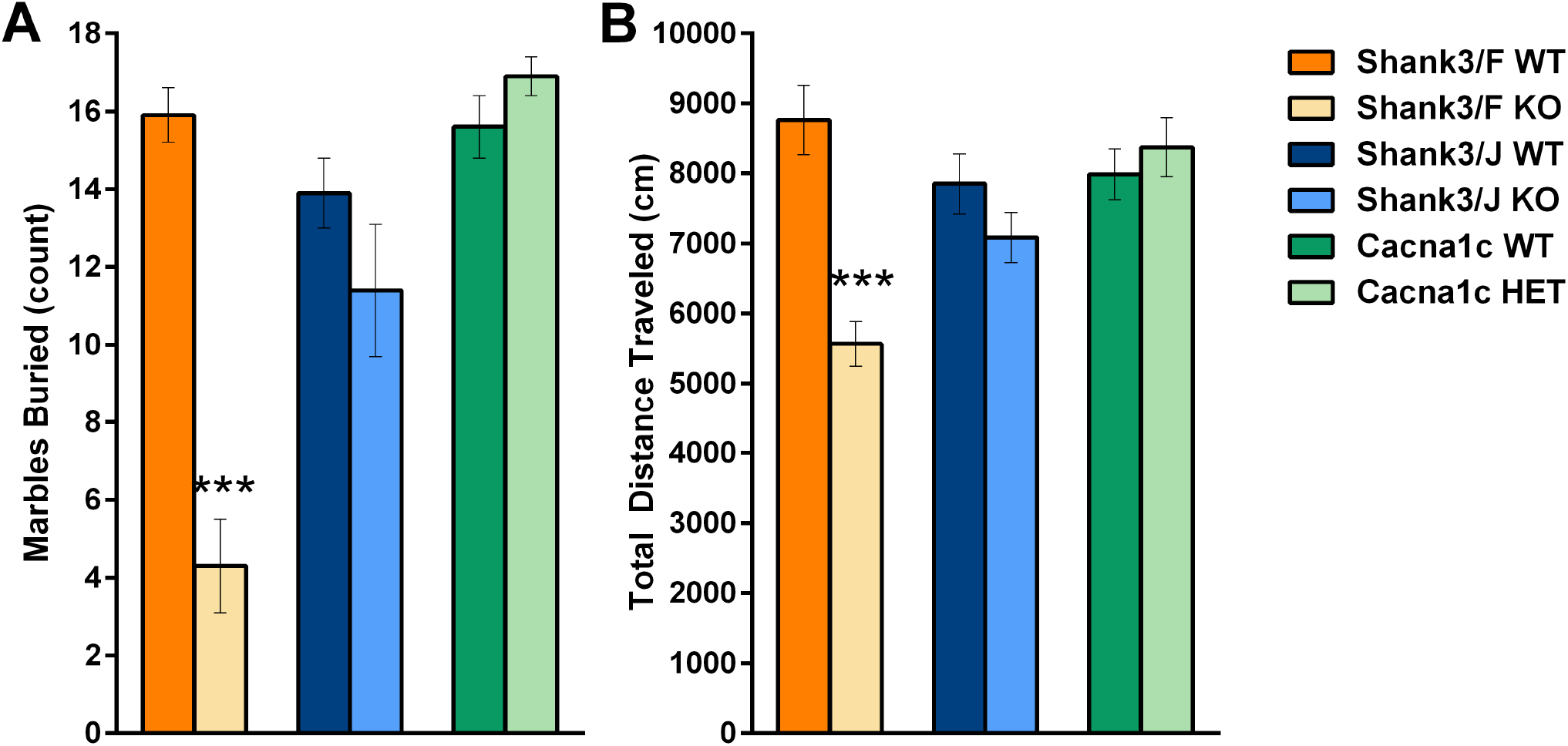
In the marble burying test, marbles are placed on the surface of a novel diggable medium. A: Number of marbles buried was decreased for Shank3/F but not Shank3/J or Cacna1c mutant mice compared to their respective WT controls. B: Locomotor activity in Shank3/F mutant mice was lower than that WT control mice, whereas the Shank3/J and Cacna1c mutant mice were as active as the corresponding WT mice. Data shown are means ± SEM; n=16 mice per genotype/line. (***p< 0.001)

Because the report by Peça et al. [7] described dramatic skin lesions, we assessed 64 and 36 *Shank3/F* mice from the colony at 4.5 and 6.5 months of age, respectively (equal numbers of WT and KO). At 4.5 months of age, we found minor balding patches in 4 WT and 7 KO mice with no lesions or signs of inflammation. At 6.5 months of age there were 3 WT and 3 KO mice with minor balding and 2 KO mice with lesions. Therefore, our finding are consistent yet milder than those published, suggesting some causative environmental factor for the lesions in Peça’s study. For *Shank3/J* and Cacna1c, we did not see increased grooming. At 4.5 months, we observed two WT mice (out of 64) and one KO mouse (out of 28) of the *Shank3/J* mice with minor balding patches. No lesions were seen in either model at this age or at 6.5 months. We did not observe the reported handling seizures in *Shank3/F* KO mice.

## Summary and Discussion

As part of a larger project to further characterize five genetic models of autism, here we have presented the data obtained from two distinct mouse models of *Shank3* mutations and heterozygous mutation in Cacna1c. We used a comprehensive behavioral battery covering multiple functional domains, including but not restricted to core autism-like features.

Starting at P4, we assessed behavior, health, and developmental milestones. Our results reproduce some of the published results for *Shank3/F* KO mice as described below. *Shank3/J* KO mice showed a milder phenotype overall compared to *Shank3/F* KO mice and, as previously shown [8], were healthy and showed no major abnormalities. Body weight was not increased at the ages investigated here, in contrast with the initial published report [8]. Pup vocalizations were low in general in both *Shank3* models with our protocol, and thus probably inconclusive due to a floor effect. We saw no other phenotypic difference in the neonatal tests. Cacna1c HET mice showed a mild phenotype overall, were healthy and showed no major abnormalities. The number of pup vocalizations for both *Shank3* mutant mice and controls increased similarly with age, with the greatest number emitted at P15, a pattern we have described in other C57 pups [1]. However, it is worth noting that pup vocalizations were low in general in both *Shank3* models with our protocol, and thus probably inconclusive due to a floor effect.

We ran two novel tests that provided comprehensive and unbiased phenotyping of activity levels in the models. In these tests we observed minor yet significant differences in gait and activity. Both *Shank3* KO models showed a decrease in activity, although this was less pronounced in the *Shank3/J* than in the *Shank3/F* model, it is consistent with the published report [8]. SmartCube did reproduce the published findings of reduced rearing in *Shank3/F* KO mice. For the Cacna1c model, we observed no significant differences in gait but did see a decrease in activity in several tests. Cacna1c HET mice showed reduced locomotion in the reciprocal social interaction and, consistent with the published report [10], traveled less distance during the baseline session in the urine open field. Neurocube, our gait analysis platform, showed mild differences in *Shank3/F* KO and *Shank3/J* KO mice and no differences in Cacna1c HET mice, when compared to their controls, consistent with published reports of normal motor abilities and reflexes [10].

Repetitive and anxiety-like behavior was assessed in the marble-burying test. In this test, *Shank3/F* KO mice buried fewer marbles than control mice, which can be interpreted as a result of reduced perseveration and anxiety-like behavior, but it may also be a secondary consequence of the general hypoactivity observed in these mice. There were no differences across genotypes for the *Shank3/J*. There also were no differences across genotype for the Cacna1c HET model, contrary to reports that Cacna1c HET mice buried twice the number of marbles [10].

Peca et al. [8] reported significant overgrooming and striking skin lesions in the *Shank3/F* KO mice. Consistent with this, SmartCube also picked up increased grooming in *Shank3/F* KO mice as a top behavioral feature. However, it is notable that the incidence of skin lesions was markedly less than the previous report. The lack of reproducibility of this extreme phenotype highlights the importance of replication studies such as ours. In discussing possible factors contributing to to this discrepancy, it is likely that differences in housing conditions may have contributed to the differing results (Dr. Feng, personal communication).

When we analyzed social behaviors, we observed differences in the urine exposure open field, a, test not previously assessed in these models. However, we failed to find robust social deficits in any of the mutant models in the three-chamber social interaction and in the reciprocal social interaction tests, with some caveats as described below.

Statistically, *Shank3/F* KO mice showed no preference for the chamber with a social stimulus during the sociability phase; however, this cannot be interpreted as a social deficit per se because WT controls similarly lacked a preference for the social stimulus chamber. (Note: we used two-factor ANOVAs to look for significant differences in differential exploration, whereas many papers use a single-factor ANOVA that represents a more liberal statistical test.) During the recognition phase, there was no genotype effect in chamber preference, unlike the published report [7]. During the sociability and recognition phases *Shank3/F* KO mice sniffed the social and novel social stimulus more than the alternatives, and a reduction in the magnitude of the preference between the two genotypes did not reach significance. As in our habituation session, *Shank3/F* KO mice preferred the center chamber more than the WT mice, it is possible that the observed differences are due to a generalized anxiety-like reaction to manipulation and novelty, rather than a specific social phenotype. In the reciprocal interaction test, *Shank3/F* KO mice initiated social events as frequently as did the WT mice but remain in contact longer, contrary to the published report [7]. This difference may be due to their hypoactivity in this and other tests.

Contrary to the published results, which described a lack of preference for the social stimulus in the *Shank3/J* model [8], we did not find any genotypic differences in the three-chamber test. In the published report, KO mice showed reduced mild social investigations and spent more time grooming and sifting through the bedding materials than control C3H mice. In addition to other methodological differences (i.e., in the published report mice were isolated for 14 days prior to testing), we purposely ran this test without bedding or marking of mice to minimize the chances of enhanced stimuli-driven behavior that could reduce social behavior. During exposure to the urine in the urine open field test, *Shank3/J* KO mice vocalized less than wild type mice. This is unlike the published result of increased vocalizations in the male mice in a reciprocal interaction test.

In the three-chamber test, no differences in preference for the chamber with a social stimulus during the sociability phase was observed between Cacna1c HET mutant mice and WT controls, similar to the published report [10]. In the reciprocal interaction test, Cacna1c HET mice initiated social events as frequently as did the WT mice but remained in contact slightly longer, and spent more time in close proximity, which may be due to their hypoactivity in this and other tests.

For both *Shank3* models, in both sessions of the urine open field test, we observed increased avoidance of the center of the arena, resembling the published report of reduced open arm exploration in the elevated plus maze and longer latencies to enter the lit area in a light-dark test [7]. In the urine open field test, during exposure to the urine of estrous females, the *Shank3F and Shank3/J* KO mice vocalized less than WT mice. We observed avoidance of the center of the arena during the exposure session but not during the baseline session in the Cacna1c HET mice. This could be due to increased anxiety or less attraction for the social stimulus. As the urine-open field test is run in low light conditions and with home bedding placed in a corner, explanations other than anxiety need be considered (i.e. increased thigmotaxis). This is in contrast with the lack of anxiety-like phenotype in the *Shank3/J* model in the published report [8]. Likewise, in the published report, Cacna1c HET did not show increased thigmotaxis, nor did they show not increased anxiety [10].

We found no cognitive or performance deficits a procedural T-maze during the acquisition or reversal phase for any of the models. This is consistent with published lack of deficits in the Morris Water Maze for the *Shank3/F* [7] but contrary to the published deficits in the novel object recognition and mild deficit in reversal in the Morris water maze [8]. As the T-maze test is expected to rely heavily on dorsolateral and dorsomedial striatal function [35], it was surprising not to see deficits in the *Shank3/F* given their reported electrophysiological striatal deficits [7].

Startle and sensory motor gating (PPI) are an important reflex and pre-attentive responses, [31]. *Shank3/F* mice showed a reduced startle response and increased prepulse inhibition of startle, findings not expected in particular for ASD since they are not domains typically of focus in the study of ASD. This was a positive, novel result not explored in the original study [7]. Consistent with the published results, startle response and prepulse inhibition of startle showed no genotype effects for the *Shank3/J* or Cacna1c models [8, 10].

We would like to emphasize that the overarching goal of this large phenotyping study was not to replicate all protocols used in previous reports. Instead, our goal was to detect robust phenotypic differences in ASD-relevant behavioral domains and to perform the same tests in the same manner to allow for comparison across several important models. Thus, robustness of the phenotype, i.e. persistence of the phenotypic deficit across small protocol variations, rather than exact replication was the central aim of this project. Deviations in specific methodologies, laboratories, and enrichment conditions present a challenge for all researchers and provide an explanation for the appearance of phenotypes milder than those reported elsewhere. Many of the current tests for ASD mouse models, especially those with a social component, appear to be particularly sensitive to procedural variables and are often confounded with activity. The genetic background can also be an explanation for differences found in our current study compared to previous ones. In the originally published studies, *SHANK3/F* and Cacna1c were on mixed backgrounds, whereas both *Shank3* lines and the Cacna1c line now available at The Jackson Laboratory, and used for the current study, are fully congenic for C57B6J.

The absence or dysfunction of SHANK3 has been associated with Phelan-McDermid syndrome (PMS), in which a portion of the chromosome 22 is missing. Individuals with PMS present delayed speech and development, decreased sensitivity to pain, unsteady ataxic gait, hypotonia, poor thermoregulation, craniofacial and hand abnormalities, seizures and locomotor deficits [36]. Hypotonia may resemble the hypoactive profile found in these *Shank3* mouse models, although individuals with PMS present with hyper rather than hypoactivity. Ten percent of the individuals with PMS show increased weight gain [37], similar to what we observed in *Shank3/F* and was previously reported in *Shank3/J* [8]. Gait in PMS is characterized by a progressive rigidity of the posture with shuffling gait and broad base [37, 38]. Although rigidity could relate to our findings of reduced variability in paw position, a shuffling gait, associated with reduced speed, and an accompanying broad base seem opposite to what we describe in this paper. Although anxiety does not seem to be a cardinal feature of the disorder it may be present (Phelan & Rogers, 2011) consistent with some aspects of the phenotypes describe here.

Mutations in the Cacna1c HET mice are homologous to those found to underlie Timothy syndrome [9]. Individuals with Timothy syndrome have a very short lifespan (2.5 y average) likely due to heart abnormalities. As it is a very rare disorder with such high mortality, a detailed description of the behavioral and cognitive symptoms is not available. However, for a model with construct validity, abnormal functional readouts can be used effectively for development of novel therapeutics, without the need for extensive and exact mapping between the mouse and the human symptoms.

Attributing rodent behavioral deficits to domain categories can be useful to organize and understand potential suites of behavioral repertoires that may correspond to human pathology. However, abnormal functional readouts not easily mapped to the human condition can also be exploited in the rodent to assess the effectiveness of novel therapeutics.

In summary, we have shown that the *Shank3/F* KO mice have a robust phenotype suggesting increased anxiety-like features, particularly in response to manipulation, novelty or salient stimulation with putative social behavior deficits. *Shank3/J* KO mice, on the other hand, showed a milder but somehow similar phenotype with some deficits in social behavior, and possibly increased anxiety-like reactivity. Both *Shank3* models showed similar mild motor behavior characteristics, namely, a narrower and more rigid gait. The Cacna1c HET mice showed a mild hypoactive phenotype without robust deficits in social behavior. We suggest that other, more exhaustive cognitive tests tackling behavioral flexibility may be of interest to further investigate cognitive deficits in the Cacna1c mouse model.

We believe that, in the extent that the genetic insult (etiological validity) [39] recapitulates the human condition, models are useful to understand gene function and pathophysiology and, therefore, to assess putative treatments. Our challenge is to pick endpoint measures as well as physiological endpoints and biomarkers that are affected by the pathological process in rodents and human in a homologous way (construct validity), and that present maximal robustness and replicability. Relying purely on analogy (i.e. face validity) has not been a fruitful approach, especially in the ASD preclinical realm.

## Materials and Methods

### Ethics Statement

PsychoGenics is an AAALAC accredited facility (Unit Number – 001213) and work is performed under PHS OLAW Assurance #A4471-01. This study was carried out in strict accordance with the recommendations in the Guide for the Care and Use of Laboratory Animals of the National Institutes of Health. The protocol was approved by the Committee on the Ethics of Animal Experiments of PsychoGenics. All efforts were made to minimize suffering and maximize animal welfare.

### Subjects

#### *Shank3* Feng line

Breeders: A cohort of 40 Het female and 20 male mice (JAX 3258240; stock 017688, B6.129-*Shank3* <tm2Gfng>/J) was provided by The Jackson Laboratory at 7.4-10.0 weeks of age. Line development prior to arrival at JAX: Exons 13-16 of SH3/ankyrin domain 3 were replaced with a neomycin resistance (neo) cassette. The construct was electroporated into (129X1/SvJ X 129S1/Sv)F1-Kitl^+^-derived R1 embryonic stem (ES) cells. Correctly targeted ES cells were injected into C57BL/6 blastocysts and the resulting chimeric males were bred to C57BL/6J females. The offspring were then intercrossed for five generations and maintained on a mixed C57BL/6J X 129 background prior to sending to The Jackson Laboratory. Line maintenance at JAX: Upon arrival, mice were additionally backcrossed to C57BL/6J inbred mice (Stock No. 000664) using a marker-assisted, speed congenic approach to establish this congenic line. Genome Scan results indicated that *Shank3* Feng breeders were fully congenic for C57B6J.

#### *Shank3* Jiang line

Breeders: A cohort of 40 Het female and 20 male mice (JAX 3258237; stock 017442, B6.129S7-*Shank3* <tm1Yhj>/J), was provided by The Jackson Laboratory at 5.6-13.1 weeks of age. Line development prior to arrival at JAX: To create this model, exons 4-9 of the SH3/ankyrin domain 3 were replaced with a neo cassette into a 129S7/SvEvBrd-Hprtb-m2-derived AB2.2 ES cells. Correctly targeted ES cells were then injected into C57BL/6J blastocysts and the resulting chimeric males were bred to C57BL/6 females. These mice were backcrossed for at least 7 generations to the C57BL/6J background. The Disc1 mutation in the 129SvEv ES cells was bred out. Line maintenance at JAX: Upon arrival at The Jackson Laboratory, mice were bred to C57BL/6J (Stock No. 000664) for at least one generation before being shipped to PsychoGenics. Genome Scan results indicated that *Shank3* Jiang breeders were fully congenic for C57B6J.

#### Cacna1c line

Breeders: A cohort of 40 WT female and 20 Het male mice (JAX 3474516 & 3490636; stock 019547, B6.Cg-Cacna1ctm2Itl/J) was provided by The Jackson Laboratory at 6.0-15.0 weeks of age. Line development prior to arrival at JAX: The TS2-neo construct designed by Dr. Rasmusson (SUNY Buffalo) was introduced in exon 8. TS2-neo mice were backcrossed with C57BL/6J for at least nine generations prior to sending to The Jackson Laboratory Repository in 2013. Line maintenance at JAX: TS2-neo mice were rederived and further backcrossed on a C57BL/6J background.

### PsychoGenics breeding scheme

Mice were set in trios (2 females:1 male) and left together for three days. Breeding was done twice to generate experimental animals. Breeders were 9.1-11.7 and 16.3-18.9 weeks of age when bred for the *Shank3/F* line cohorts 1 and 2, 7.3-14.9 and 14.4-22 weeks of age when bred for the *Shank3/J* line cohorts 1 and 2 and 8.0-17.0 and 15.0-25.0 weeks of age when bred for the Cacna1c line cohorts 1 and 2. See Table 5 in S1 Methods for breeding efficacy, gender, and genotype ratios. Sex and genotype ratios were unbiased in all models. The first cohort was weaned at 3 weeks of age in preparation for P30 testing. Animals in cohort 2 were weaned at 4 weeks of age due to small body size. Breeding success and pup survival was similar among the four groups. All testing was done in male mice. Mutants and their wild type controls were littermates. As specified by the original developer of the test [40], two-month old 129SVE male mice (Taconic Farms) were used as stimulus mice in the three-chamber test. Age-matched male (∼P45) unfamiliar littermates were used as stimulus mice for the reciprocal social interaction test.

## General Procedures

Mice were tested singly in all tests except for those that employ stimulus mice (the Reciprocal Interaction and Three-Chamber tests). For all information regarding general procedures and testing protocols, bioinformatics, and data handling, please refer to [1] and S1 Methods.

## Acknowledgments

We thank Guoping Feng, Yong-hui Jiang, and Richard Tsien for scientific discussion and feedback on an earlier version of this manuscript. We are grateful to Judy Watson-Johnson, Vanessa Rivera, and Melinda Ruiz for breeding efforts. Matt Mazzella, Andy Farrar, and others helped with quality control of data, graphs, and analysis. We are thankful for their efforts. Thanks to Igor Filippov for assistance with all video-based systems and to Mitch Silverstein who offered invaluable support in apparatus development.

## Supporting Information

**S1 Methods. PCR assays, time-course of neonatal tests, breeding output, and bioinformatics for SmartCube and NeuroCube.**

**S1 Table. SmartCube results for the three models.**

**S2 Table. NeuroCube results for the three models.**

**S3 Table. General health data for the three models.**

**S4 Table. Activity and ultrasonic vocalizations in the three models.**

**S5 Table. Motor coordination and reflexes in the three models.**

**S6 Table. 3-chamber test for the three models.**

**S7 Table. Reciprocal social interaction test for the three models.**

**S8 Table. Urine exposure open field test for the three models.**

**S9 Table. T-maze test for the three models.**

**S10 Table. Startle and prepulse inhibition of startle for the three models.**

**S11 Table. Marble burying test for the three models.**

**S12 Table. Neonatal testing raw data.**

**S13 Table. 3-chamber test raw data.**

**S14 Table. Reciprocal social interaction test raw data.**

**S15 Table. Urine exposure open field test raw data.**

**S16 Table. T-maze test raw data.**

**S17 Table. Startle and prepulse inhibition of startle raw data.**

**S18 Table. Marble burying test raw data.**

**S1 Fig. Neonatal tests.** During the neonatal, at the three ages studied, mutants and their corresponding controls did not show phenotypic differences for A) the number of grid-paper squares crossed and B) baseline temperature. Data shown are means ± SEM.

**S2 Fig. The three-chamber test of social behavior showed minor differences between mutants and the corresponding WT mice.** A: During the baseline habituation session SHANK3/F KO mice showed less exploration of the side chambers than the corresponding WT mice. In the Cacna1c model, both genotypes showed more exploration of the right side chamber than the left side chamber. B-C: A mouse preference sociability (B) and recognition (C) index built combining the time in the side chambers showed no genotypic differences for any of the models. D: During the sociability test, whereas for the SHANK3/F and Cacna1c models there were no effects of genotype or chamber type, SHANK3/J mice and their control littermates crossed over to the mouse chamber more than to the object chamber. E: During the recognition test, the SHANK3/F KO mice moved across chambers more frequently than wildtype controls and there was an overall increased crossing to the novel mouse chamber for both models. Cacna1c mutant mice moved across chambers less frequently than wildtype controls. Data shown are means ± SEM. Asterisks refer to differences between genotypes. Numerals refer to differences between chamber types (Chamber side main effect: ^##^p< .01, ^###^p< .001; Genotype main effect: *p< .05).

**S3 Fig. In the reciprocal interaction test, where pairs of mutant or wild type mice are allowed to interact in an arena for 10 minutes (see [1] for complete methods), there were differences in activity.** Mutant mice travelled less distance compared to WT controls. Data shown are means ± SEM (*p< .05, **p< .01, ***p< .001).

**S4 Fig. The percent of the 10-minute session time that pairs of mutant or wild type mice spent interacting with each other was increased only on the reciprocal interaction measure in SHANK3/F mutant mice compared to WT.** Active interaction is where the subject mouse explores the stimulus mouse, passive interaction is where the stimulus mouse explores the subject mouse, and reciprocal interaction is where the subject and stimulus mouse explore each other (see [1] for complete methods). Data shown are means ± SEM (*p< .05).

**S5 Fig. In the procedural T-maze, where mice are trained to enter one side of the maze to reach a platform and escape the water, mutant and WT mice learned the task at a similar rate as shown by the percent correct choice during reversal, when mice are required to go to the opposite side of the maze to reach the platform (see [1] for complete methods).** Data shown are means ± SE.

